# The Eyes Absent family members EYA4 and EYA1 promote PLK1 activation and successful mitosis through tyrosine dephosphorylation

**DOI:** 10.1101/2022.10.10.511510

**Authors:** Christopher B. Nelson, Samuel Rogers, Kaushik Roychoudhury, Yaw Sing Tan, Caroline J. Atkinson, Alexander P. Sobinoff, Christopher G. Tomlinson, Anton Hsu, Robert Lu, Eloise Dray, Michelle Haber, Jamie I. Fletcher, Anthony J. Cesare, Rashmi S. Hegde, Hilda A. Pickett

## Abstract

The Eyes Absent family of proteins (EYA1-4) are a biochemically unique group of tyrosine phosphatases known to be tumour promoting across a range of cancer types. To date, the molecular targets of EYA phosphatase activity remain largely uncharacterised. Here, we identify Polo-like kinase 1 (PLK1) as a direct interactor and phosphatase substrate of both EYA4 and EYA1, with pY445 on PLK1 being the primary target site. EYA-mediated dephosphorylation of PLK1 in the G2 phase of the cell cycle is required for centrosome maturation, PLK1 localization to centrosomes, and polo-box domain (PBD) dependent interactions between PLK1 and the PLK1-activating proteins BORA and CEP192. Molecular dynamics simulations support the rationale that pY445 confers a structural impairment to PBD-substrate interactions that is relieved by EYA-mediated dephosphorylation. Depletion of EYA4 or EYA1, or chemical inhibition of EYA phosphatase activity, dramatically reduces PLK1 activation, causing mitotic defects and cell death. Overall, we have characterized a novel phosphotyrosine signalling network governing PLK1 and mitosis. This work provides a mechanism of cell killing for EYA phosphatase inhibitors with important therapeutic implications.

## Introduction

The Eyes Absent Family (EYA1-4) are a unique group of dual-function proteins with oncogenic roles in a variety of tumour types, where they promote cancer cell phenotypes such as proliferation, survival and migration ^1–9^. EYA proteins possess N-terminal transcriptional coactivation activity and C-terminal Haloacid Dehydrogenase (HAD) protein tyrosine phosphatase activity ^5, 10–19^. EYAs are the only known HAD-family tyrosine phosphatases and have unique active site chemistry making them well suited to specific inhibition with small molecules^20–23^. This has led to the identification of EYA phosphatase inhibitors, and there is growing interest in their chemotherapeutic potential ^20, 24–27^. However, the molecular functions and specific targets of EYA phosphatase activity are largely unknown. This is particularly the case for EYA4, which currently has no known phosphatase substrates.

Here, we employ an unbiased proteomics approach to identify EYA4 substrates that both interact with EYA4 and have tyrosine phosphorylation levels that respond to genetic or pharmacological perturbation of EYA4. We identify the master mitotic regulator PLK1, and specifically Y445, a residue within polo-box 1 of PLK1, as a *bona-fide* substrate of both EYA4 and EYA1. EYA4 and PLK1 interact and colocalize specifically at centrosomes in G2 cells, and this interaction is dependent on a conserved polo-docking site (PDS) present on EYA4 and EYA1. We demonstrate that dephosphorylation of pY445 by EYA4 and/or EYA1 promotes PLK1 localization to centrosomes and centrosome maturation, normal spindle morphology, PLK1 PBD dependent interactions, and PLK1 kinase activation. This mechanism is supported by structural simulations, which predict that Y445 phosphorylation reduces substrate recognition and alters polo-box function through flexibility changes within the connecting loop. Finally, we show that EYA-mediated dephosphorylation of PLK1 supports PLK1 function during mitotic progression, while treatment with an EYA phosphatase inhibitor potently elicits cell death in tumour cells expressing EYA4 and/or EYA1.

## Results

### EYA4 interacts with and dephosphorylates PLK1

To identify EYA4 substrates, we fused full length EYA4 to a Myc-tagged BioID2 biotin ligase and identified EYA4 interacting proteins using a BioID2 proximity proteomics strategy ^28, 29^ (Figure 1A-B, S1A-C). Biotinylation of proteins after expression of Myc-BioID2-EYA4 was compared to a BioID2-Myc control using label free quantification (LFQ). This yielded 156 high confidence EYA4 interactors (>4-fold enrichment, p-value <0.05) (Figure 1A, Table S1). Polo-like kinase 1 (PLK1) and Aurora kinase A (AURKA), two of the master regulatory kinases that control mitosis, were among the most highly significant EYA4 interacting proteins (p < 0.00001 and 0.0001 respectively, Figure 1A, Table S1). PLK1 and AURKA localize to mitotic structures including the centrosome and spindle midbody, with PLK1 also localizing to the spindle and kinetochore^30^. To determine whether EYA4 is likely to share cell cycle specific localization with PLK1 and AURKA, we mined the EYA4 interactome for other centrosomal or spindle proteins. We identified 54 EYA4 interactors with functional relationships to the centrosome, spindle, or mitosis. Additionally, 24 of these proteins have established localization to mitotic structures, including 17 with centrosomal localization (Figure 1B, Table S1).

**Fig. 1.**
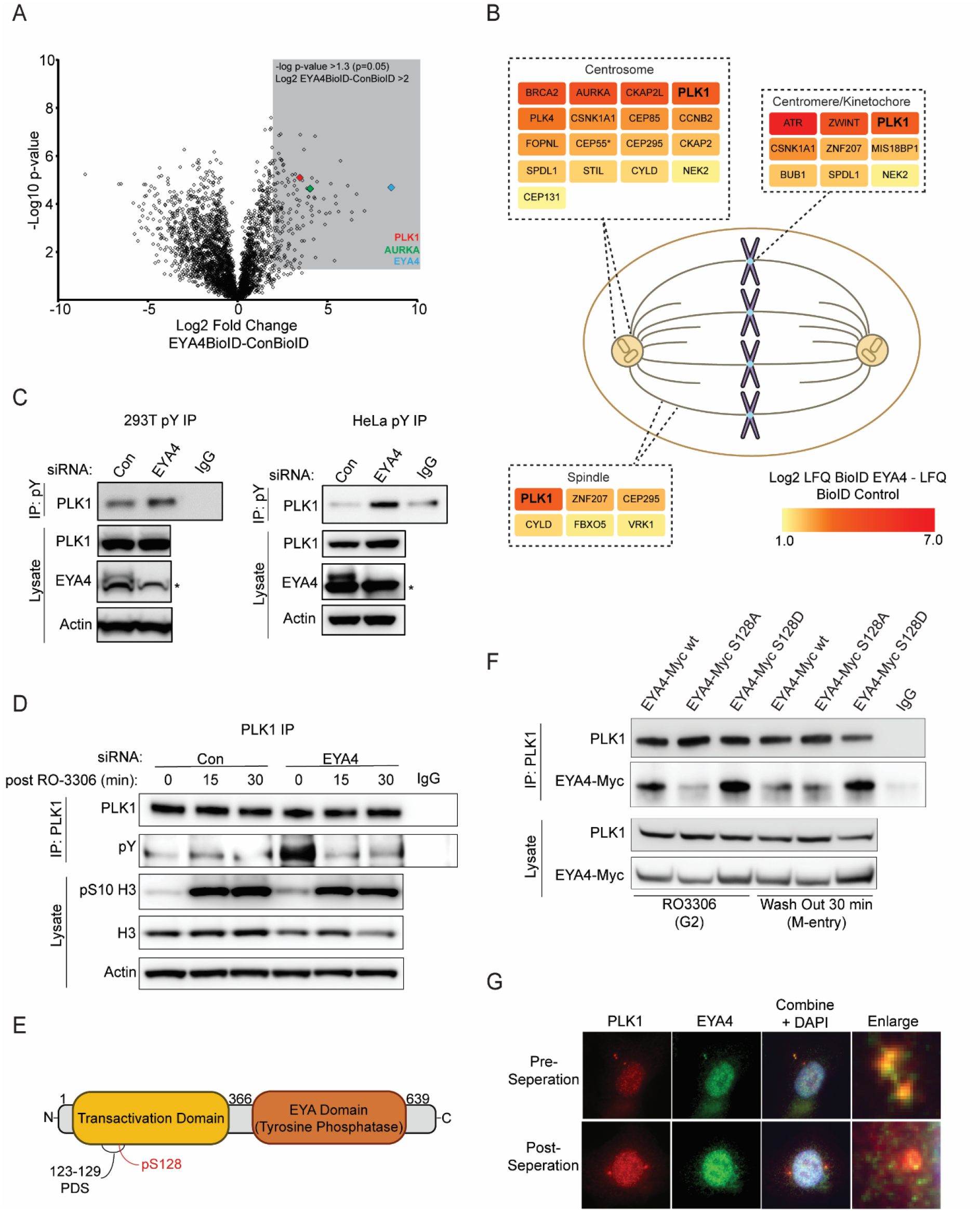
**a,** Volcano plot of EYA4-Myc-BioID interactome (EYA4BioID, right side) vs Myc-BioID interactome (ConBioID, left side). Grey coloured box represents 156 high confidence EYA4 interactors with -log p-value > 1.3 and Log2 fold difference > 2. EYA4, PLK1 and AURKA are indicated on the plot (n=4). **b,** EYA4 interactors with known localization to mitotic structures including PLK1. Proteins have been colour coded by Log2 fold difference. **c,** Tyrosine phosphorylated proteins were immunopurified (IP) from 293T or HeLa protein lysates. Asterisk represents non-specific band on the EYA4 blot (n=3). **d,** Immunopurified endogenous PLK1 from control or EYA4 depleted 293T cells that had been arrested in G2 with RO3306 or released into mitosis for 15 or 30 minutes and immunoblots performed using an anti-phosphotyrosine antibody (pY). **e,** Schematic representation of EYA4 protein with N-terminal transactivation domain and C-terminal tyrosine phosphatase domain as well as putative PDS (AA 123- 129) and known phosphosite (pS128). **f,** Co-immunopurification reactions between immunopurified endogenous PLK1 and an EYA4-Myc construct or EYA4-Myc mutants. Cells have been arrested in G2 with RO3306 or released into early M phase. **g,** Immunoflourescence of endogenous PLK1 and EYA4 in G2 arrested cells. A representative image is shown of a cell that is in the process of centrosome separation (pre-separation) as well as a cell that has fully separated centrosomes (post-separation).

To determine candidate EYA4 tyrosine phosphatase substrates, we depleted cells of EYA4 and performed immunoprecipitations using an anti-phosphotyrosine antibody followed by LC-MS/MS analysis. We identified 50 proteins with statistically significant increases in tyrosine phosphorylation, including PLK1 (p < 0.05, Figure S1D-E, Table S1). Western blots of anti-phosphotyrosine immunoprecipitations confirmed an increase in PLK1 tyrosine phosphorylation in both 293T and HeLa cells following EYA4 depletion (Figure 1C). Using a reversed experimental design, we immunopurified endogenous PLK1 from cells arrested in G2 when PLK1 is concentrated at centrosomes, or following release into M-phase, when PLK1 becomes primarily localized to kinetochores, and blotted for tyrosine phosphorylation^31^. EYA4 depletion resulted in a striking increase in PLK1 tyrosine phosphorylation in G2, but this increase did not persist into mitosis (Figure 1D). This suggests that EYA4 may regulate PLK1 function at G2 centrosomes through dephosphorylation.

### EYA4 and EYA1 contain a polo-docking site (PDS) that confers interaction with PLK1 in G2

Most PLK1 interactors possess a consensus PDS motif containing an internal phosphorylation site ^32, 33^. We identified a potential conserved PDS within the N-terminal transactivation domains of both EYA4 and EYA1, matching 6 of 7 residues of the consensus PDS sequence (AA 123-129, Figure 1E, S1F). Phosphorylation of pS128, which falls within the PDS of EYA4, has previously been observed in a high throughput dataset, suggesting that the PDS is functional (Figure 1E)^34^. We used co-immunoprecipitation to determine whether the putative PDS on EYA4, and specifically phosphorylation of S128, mediated the interaction with PLK1. Myc-tagged WT EYA4 strongly interacted with PLK1 in G2 arrested cells. This interaction was abrogated for EYA4 containing a S128A non-phosphorylatable mutation and enhanced for EYA4 with a S128D phosphomimetic mutation (Figure 1F). Further, while the WT EYA4-PLK1 interaction was lost upon entry into mitosis, the S128D-PLK1 interaction persisted (Figure 1F). These results suggest that phosphorylation of the EYA4 PDS regulates the G2-specific interaction between EYA4 and PLK1.

Consistent with an EYA4-PLK1 interaction in G2, we found that endogenous EYA4 colocalized with PLK1 at centrosomes in G2 (Figure 1G). Colocalization between EYA4 and PLK1 was strongest in cells prior to centrosome separation (Figure 1G). This suggests that EYA mediated dephosphorylation of PLK1 in G2 may regulate PLK1 functions in centrosome biology and early mitosis.

Myc-tagged EYA1 also interacts with PLK1 in G2 arrested cells, while EYA3, which lacks an obvious PDS, does not (Figure S1G). Further, depletion of EYA1, or both EYA4 and EYA1, also resulted in an increase in PLK1 tyrosine phosphorylation (Figure S1H), suggesting that EYA1 may function redundantly with EYA4 to dephosphorylate tyrosine residues on PLK1 during G2.

### EYA4 and EYA1 promote PLK1 activation

PLK1 kinase activation begins at centrosomes and in the cytoplasm during G2 via AURKA mediated phosphorylation of PLK1 at T210, reaching peak activation upon mitotic entry ^30, 35–38^. We assessed whether EYA4 or EYA1 can alter PLK1 levels or activation (pT210) in 293T and HeLa cells using automated quantification of fluorescence intensity in mitotic cells with positive staining for pS10 H3 (Figure 2A, S2A). Individual knockdown of EYA4 or EYA1 reduced PLK1 activation to differing magnitudes, with each knockdown reaching statistical significance in separate cell lines (Figure 2A). Total PLK1 intensity was slightly elevated following EYA1 knockdown in HeLa cells (Figure 2A). Co-depletion of EYA4 and EYA1 caused an additive decrease in PLK1 activation in both cell lines (p < 0.0001, Figure 2A, knockdown confirmation: S2B).

**Fig. 2.**
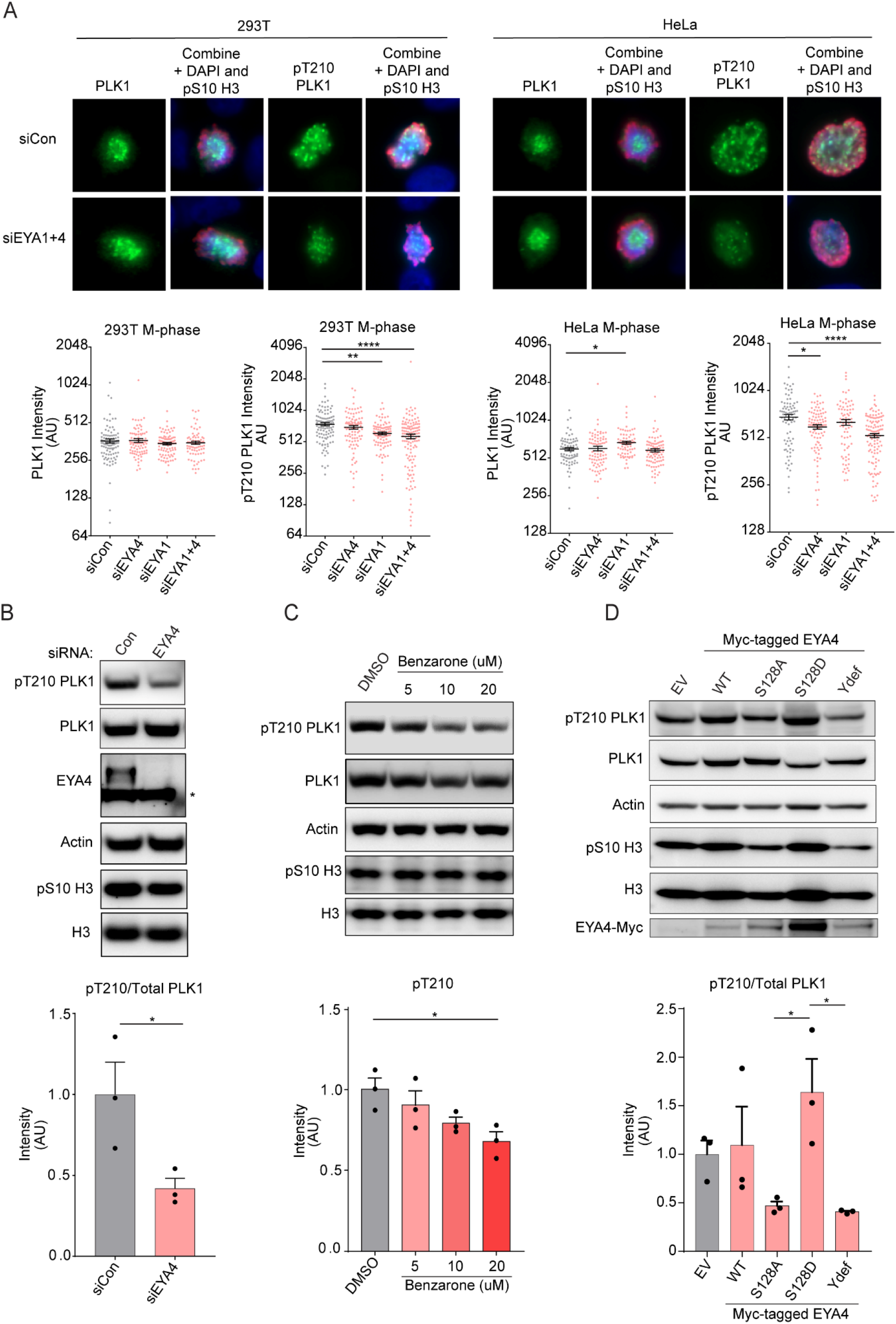
**a,** Mitotic 293T or HeLa cells were identified by positive staining for pS10 H3 in high throughput image analysis experiments. Staining and quantitation were performed for total PLK1 and pT210 PLK1 in separate experiments following knockdown of EYA1, EYA4 or the combined knockdown. Representative images from the combined knockdown experiments are shown for each cell line (top). Quantitation of fluorescence intensity of total PLK1 or pT210 PLK1 in individual mitotic cells is shown for each treatment (bottom). Combined knockdown of EYA4 and EYA1 caused a highly significant decrease in pT210 PLK1 intensity in both cell lines (p < 0.0001). **b,** Western blots from nocodazole arrested mitotic HeLa cells and densitometry of the ratio of pT210 PLK1 to total PLK1 from three biological replicates (p < 0.05, asterisk indicates a non-specific band on the EYA4 blot). **c,** Western blots following treatment with a pan-EYA phosphatase inhibitor in nocodazole arrested mitotic HeLa cells and densitometry of pT210 intensity from three biological replicates (p < 0.05 for 20μM benzarone). **d,** Representative western blots and pT210/total PLK1 densitometry in three biological replicates of nocodazole arrested mitotic HeLa cells overexpressing EYA4 or EYA4 mutants. (p < 0.05 for comparisons between EYA4 S128D and EYA4 S128D or EYA4 ydef).

Depletion of EYA4 significantly decreased pT210 levels relative to total PLK1 in western blots performed on lysates from nocodazole arrested mitotic cells (p < 0.05, Figure 2B). Further, there was a dose dependent reduction of pT210 in mitotically arrested cells treated concurrently with the EYA phosphatase inhibitor benzarone and nocodazole for 12 hours following thymidine release, reaching statistical significance in the 20 μM benzarone treatment (p < 0.05 Figure 2C, S2C).

In further support of EYA4 directly regulating PLK1 activity, overexpression of the EYA4 S128A mutant, or a D375N mutant previously shown to inactivate EYA4 phosphatase activity (herein referred to as Ydef), exerted dominant negative effects over endogenous EYA4, significantly reducing the activation of PLK1 relative to EYA4 S128D in nocodazole arrested cells (p < 0.05, Figure 2D)^24^. Lower levels of activated PLK1 were also observed following S128A mutant overexpression compared to WT EYA4 overexpression in unarrested mitotic cells. (p < 0.01, Figure S2D). These results suggest that EYA4 mediated PLK1 dephosphorylation is required for enhancing PLK1 activation by AURKA.

### EYA4 and EYA1 support centrosome maturation, separation, and PLK1 localization to centrosomes

During G2 to M transition, PLK1 activation is essential for the accumulation of pericentriolar material at centrosomes (centrosome maturation), and for the separation of mature centrosomes ^32, 39–43^. To evaluate the effects of EYA4 and EYA1 depletion on centrosome maturation and separation, we measured both centrosome number and the level of pericentrin (PCNT) accumulated in centrosomal foci in prophase cells. Individual or combined depletion of EYA4 and EYA1 led to a significant increase in the proportion of prophase cells with only one centrosome (Figure 3A-B). Prophase cells with only one centrosome also had lower centrosomal PCNT intensity following depletion of EYA4 or EYA1, with an additive effect being observed with combined depletion (Figure 3C). These results support a role for EYA4 and EYA1 in centrosome maturation and separation.

**Fig. 3.**
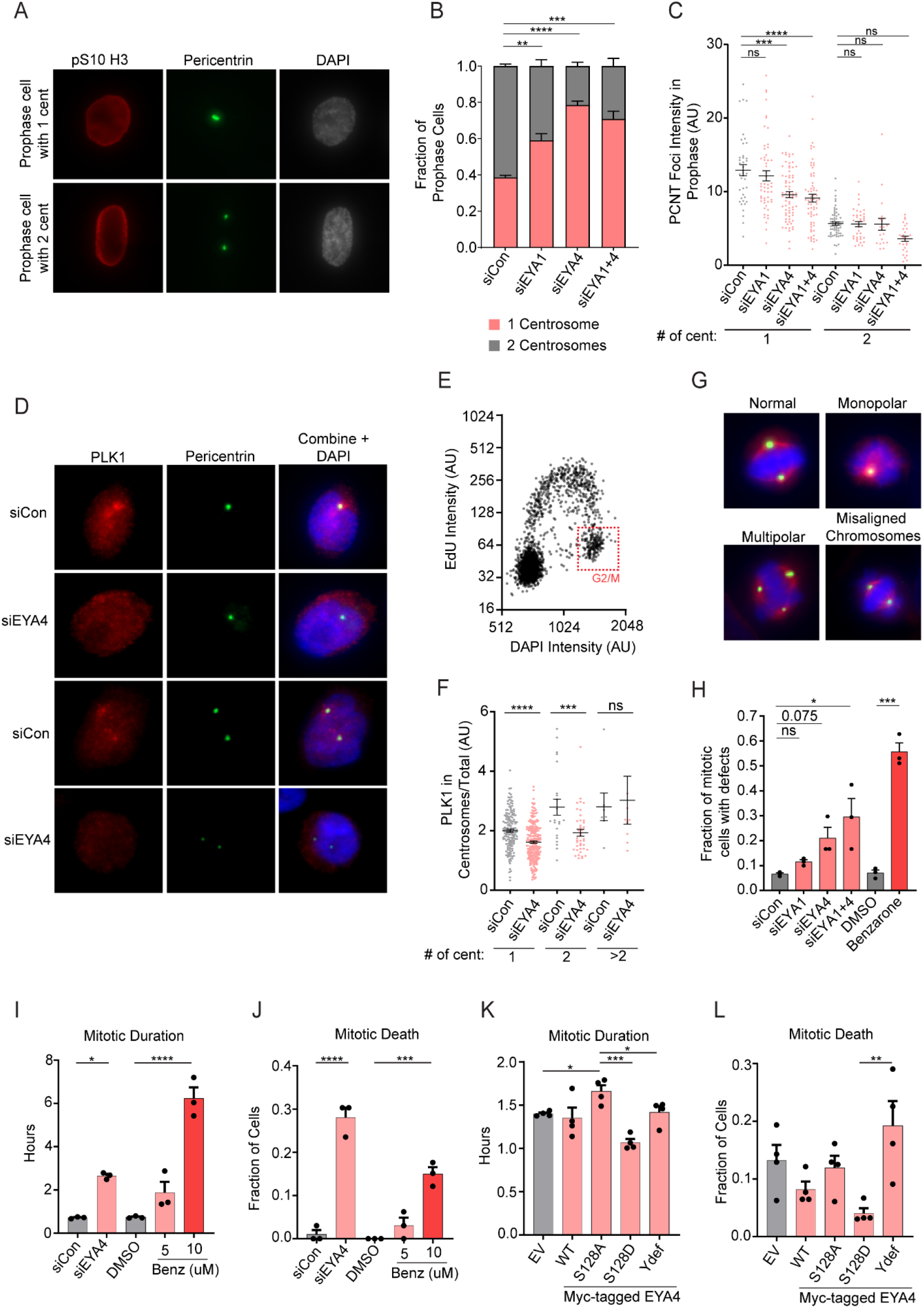
**a,** Representative images of prophase HeLa cells identified by pS10 positivity and DNA morphology with either one or two centrosomes (pericentrin foci). **b,** Quantitation of the fraction of prophase cells with one centrosome or two centrosomes following depletion of EYA1, EYA4 or the combination (p < 0.01, 0.0001, and 0.001 respectively, relative to a control siRNA, n=3). **c,** Integrated intensity of pericentrin foci in prophase cells following depletion of EYA4, EYA1 or combined EYA4 and EYA1 depletion (p < 0.001 and 0.0001 respectively in cells one centrosome, n=3). **d-e,** Asynchronous HeLa cells were pulsed with EdU for 1 hour and incorporated EdU was detected alongside PLK1, pericentrin and DAPI. **d,** Fluorescent images show examples of PLK1 colocalization with pericentrin in EdU negative G2 cells from control or EYA4 depletion conditions. **e,** G2 cells were identified by gating the cell population using EdU intensity on the Y-axis and DAPI intensity on the X-axis. The red box indicates EdU negative, DAPI high (4N DNA content) G2 cells used in the analysis **f,** Quantitation of PLK1 integrated intensity at centrosomes in relation to **d** and **e** (p < 0.0001, 0.001 respectively for comparisons made in cells with one or two centrosomes). **g,** Example images showing pericentrin and alpha-tubulin staining of mitotic HeLa cells highlighting the most commonly observed spindle defects including monopolar and multipolar spindles as well as spindles with misaligned chromosomes. **h,** Total spindle defects were enhanced following depletion of EYA4 (p < 0.075), combined depletion of EYA1 and EYA4 (p < 0.05) or following treatment of cells with benzarone (p <0.001, n=3). **i-l,** Summary data from live cell imaging experiments (n=3). **i,** Mitotic duration was increased following depletion of EYA4 (p < 0.05) and in a dose-dependent manner following treatment with benzarone (p < 0.0001 for 10μM). **j,** Mitotic cell death was increased both with EYA4 depletion (p < 0.0001) and following treatment with 10μM benzarone (p < 0.001). **k,** Overexpression of an S128A EYA4 mutant increased mitotic duration relative to EV (p < 0.05), EYA4 Ydef (p < 0.05), and EYA4 S128D (p < 0.001). **l,** Overexpression of EYA4 Ydef increased mitotic cell death relative to EYA4 S128D (p < 0.01).

PLK1 activation and centrosome maturation are intricately linked to PLK1 centrosomal localization in G2/M^30, 36, 38^. To evaluate the centrosomal localization of PLK1, asynchronous cells were pulse labelled with EdU, and G2/M cells were gated by their high DAPI intensity (equivalent to 4N DNA content) and low EdU intensity. EYA4 depletion reduced the levels of PLK1 localization to centrosomes in G2/M cells without affecting the cytoplasmic/nuclear ratio of PLK1 (Figure 3D-F, S3A).

### The phosphatase activity of EYA4 and EYA1 prevents mitotic defects and mitotic cell death

We then examined HeLa cells for mitotic spindle defects following depletion of EYA4 and EYA1 or treatment with benzarone. Total spindle defects increased modestly following EYA1 and EYA4 depletion, and significantly following combined EYA4 and EYA1 depletion (p < 0.05, Figure 3G-H). Total spindle defects were strongly induced by treatment with benzarone (p < 0.001, Figure 3G-H). Monopolar spindles and spindles with misaligned chromosomes, two defects known to be caused by PLK1 deficiency, were frequently observed following combined depletion of EYA4 and EYA1, or benzarone treatment (p < 0.001, Figure S3B)^44–47^.

We next observed the effects of EYA4 and EYA phosphatase activity on mitotic duration and mitotic cell death. EYA4 depletion prolonged mitosis and induced mitotic cell death (Figure 3I-J). These defects were also observed with benzarone treatment (Figure 3I-J). Further, benzarone caused a dose-dependent elevation in cleaved PARP, indicative of cell death, and phosphorylation of H3 on S10, indicative of mitotic arrest, in cancer cell lines with high expression levels of EYA4 (HeLa), EYA1 (SKNAS), or both (SKNFI) (Figure S4A-B).

A significant increase in mitotic duration was also detected when we overexpressed the EYA4 S128A mutant that cannot interact with PLK1, indicative of a dominant negative effect (Figure 3K). Further, overexpression of the EYA4 Ydef mutant significantly increased mitotic death compared to the S128D mutant, supporting the rationale that dysfunctional EYA4 has a dominant negative impact on PLK1 mediated mitotic progression (Figure 3I). These data demonstrate that EYA4 and EYA1 function redundantly to promote the accurate completion of mitosis through the dephosphorylation of PLK1.

### EYA4 and EYA1 target pY445 on PLK1

To investigate the mechanism by which tyrosine dephosphorylation of PLK1 by the EYAs regulates PLK1 function, we used targeted mass spectrometry to determine the specific PLK1 tyrosine phosphosites dephosphorylated by EYA4 and EYA1. Myc-tagged PLK1 was immunopurified following knockdown of EYA4 or EYA1, overexpression of EYA4 Ydef, or following benzarone treatment. Purified PLK1 was digested in-gel and subjected to mass spectrometry. Pilot experiments determined the mass and charge of four tryptic peptides containing PLK1 tyrosine phosphosites, pY217, pY421, pY425, and pY445 (Table S2). These four phosphopeptides were quantified by parallel reaction monitoring (PRM) (Figure 4A). Only pY445 peptides exhibited increased intensity relative to control across all conditions tested, making it a high confidence substrate for both EYA4 and EYA1 (Figure 4B). In addition, we observed an increase in pY425 by PRM in all conditions, except for knockdown of EYA4 (Figure 4B). This suggests that pY425 may also be a substrate of EYA1.

**Fig. 4.**
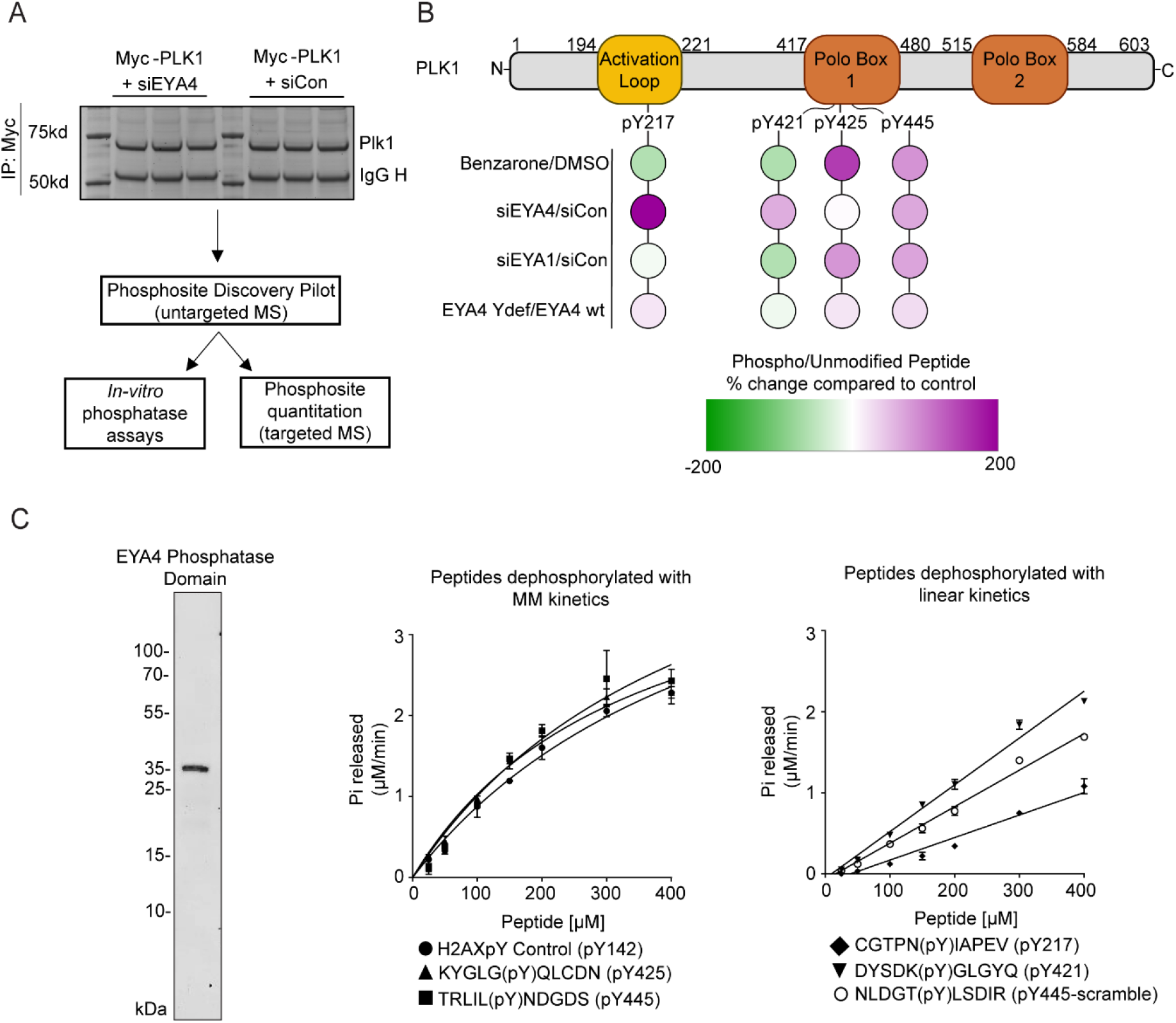
**a,** Schematic showing an example of immunopurified and Coomassie stained PLK1 which was in-gel digested and used in downstream pilot and quantitative phophoproteomic experiments. Phosphosites identified by mass spectrometry also informed in-vitro phosphatase assays. **b,** Results of quantitative phosphoproteomics experiments following a two-hour treatment with a pan-EYA phosphatase inhibitor (Benzarone, 10μM), depletion of EYA4 or EYA1, or overexpression of a phosphatase defective EYA4 protein (Ydef). Results are shown as the mean percent change in phosphorylation level relative to control treatments of four PLK1 phosphotyrosine sites (n = 2-4 biological replicates per treatment). **c,** The purified phosphatase domain of EYA4 (left, Coomassie stained SDS-PAGE gel) was used in in-vitro phosphatase assays with synthetic phosphopeptides. pY445, pY425 and a positive control peptide from H2AX (pY142) (middle), were dephosphorylated with fast Michaelis-Menten (MM) kinetics and comparable Km values (0.51mM for H2AX (pY142), 0.34 mM (pY425), 0.46mM (pY445). pY421, pY217 and a scrambled version of pY445 were dephosphorylated with slow linear kinetics (right). Error bars represent standard deviations, n=3).

To determine if EYA4 has intrinsic biochemical specificity for PLK1 phosphosites, we performed *in-vitro* phosphatase assays using the purified phosphatase domain of EYA4 (Residues 367-639) and synthetic phosphopeptides containing the four PLK1 tyrosine phosphosites (Figure 4C). A scrambled version of pY445 was used as a negative control and a peptide previously shown to be dephosphorylated by the phosphatase domains of all four EYA proteins was used as a positive control (H2AX pY142)^25, 48^. Dephosphorylation of pY445, pY425, and the positive control peptide all occurred with similar Michaelis-Menten kinetics, indicative of a specific dephosphorylation reaction (Figure 4C). In contrast, pY217, pY421 and pY445-scramble were dephosphorylated linearly with slower kinetics, suggesting a non-specific dephosphorylation reaction (Figure 4C).

To validate pY445 as the predominant phosphosite targeted by EYA4, we compared overall tyrosine phosphorylation of an unphosphorylatable Y445F mutant with WT PLK1 following EYA4 depletion in G2 arrested cells. EYA4 depletion increased overall PLK1 tyrosine phosphorylation, however no induction was observed for Y445F (Figure S5A). Therefore, pY445 is the predominant PLK1 phosphosite targeted by EYA4.

The phosphatase domains of EYA1 and EYA3 were also tested for their ability to dephosphorylate the pY445 peptide. Both EYA1 and EYA3 were able to dephosphorylate pY445 with fast Michaelis-Menten kinetics (Figure S5B). These results suggest that dephosphorylation of PLK1 pY445 is regulated both by intrinsic specificity within the highly homologous EYA phosphatase domain (~70-90% homology between EYA members), as well as by the ability of EYA4 and EYA1 to interact with PLK1 *in-vivo* through a PDS.

### A non-phosphorylatable Y445F PLK1 mutant is hyperactive

As EYA mediated dephosphorylation promotes PLK1 activity, we hypothesised that specific dephosphorylation of pY445 was responsible for this effect. In support of this hypothesis, overexpressed PLK1-Y445F had elevated pT210 phosphorylation levels relative to WT PLK1 in mitotically arrested HeLa and 293T cells (p<0.001, 0.05 respectively, Figure 5A, S5C).

**Fig. 5.**
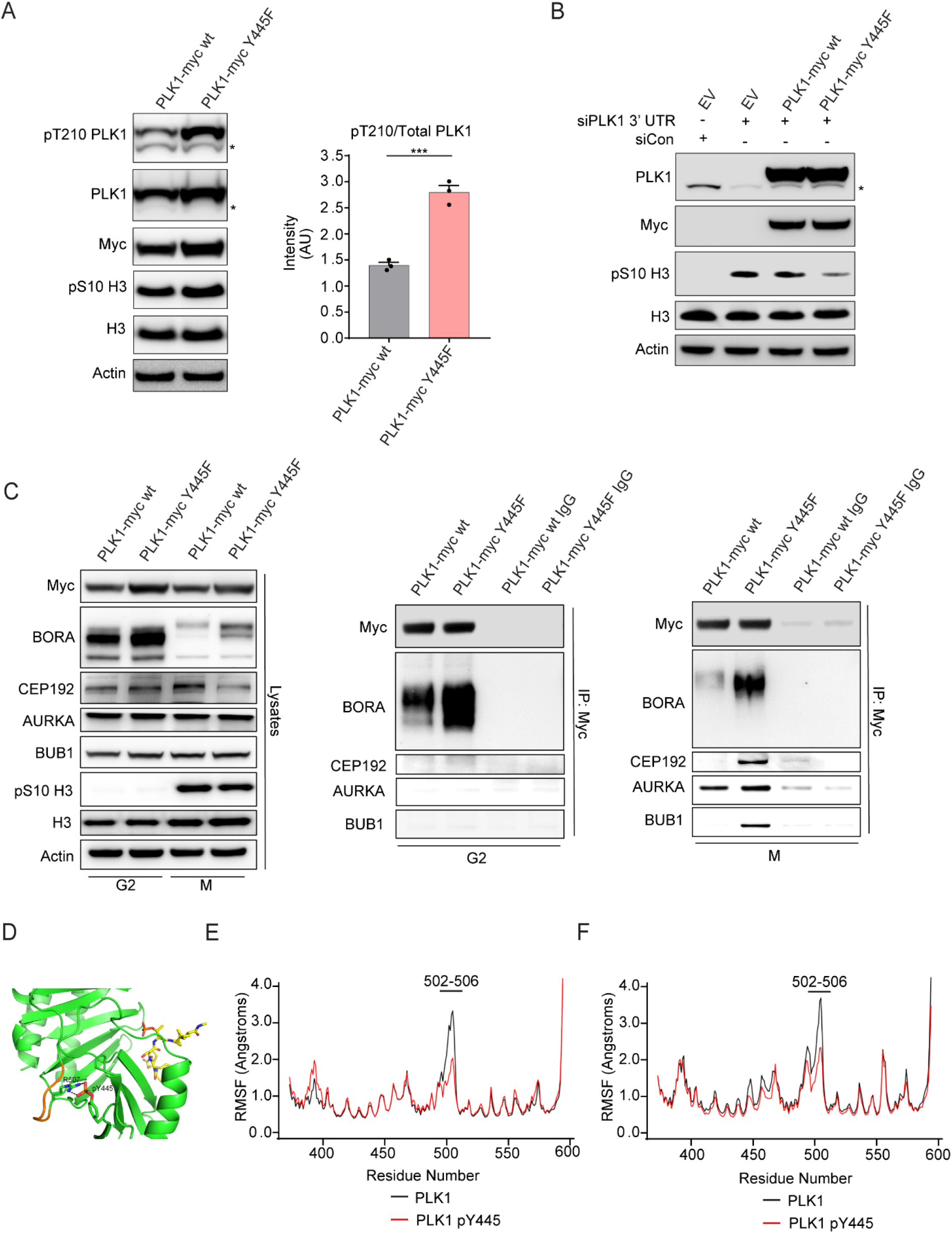
**a,** Representative western blots and pT210/total PLK1 densitometry in three biological replicates of nocodazole arrested mitotic HeLa cells overexpressing Myc-tagged PLK1 wt or a Myc-tagged Y445F PLK1 mutant (p < 0.001, n=3, asterisk on the pT210 PLK1 and PLK1 blots indicates endogenous PLK1) **b,** The Myc-tagged Y445F mutant of PLK1, but not the Myc-tagged WT PLK1 can partially rescue the mitotic arrest (pS10 H3 signal) induced by depletion of the endogenous PLK1 by a 3’UTR targeted siRNA (asterisk indicates endogenous PLK1). **c,** Coimmunoprecipitation reactions with overexpressed Myc-tagged PLK1 wt or Myc-tagged PLK1 Y445F in RO3306 arrested G2 cells and nocodazole arrested M-phase cells. **d,** A representative simulation snapshot of PLK1 PBD (green) phosphorylated at Y445 and complexed with a model phosphopeptide (yellow). The image highlights the interaction of pY445 with R507 (green sticks) and the increased rigidity within the connecting loop (AA 502-506) highlighted in orange. **e-f**, Plots of the root mean square fluctuation (RMSF) of Cα atoms in wt PLK1 PBD (black line) and Y445-phosphorylated PLK1 PBD (red line). The decreased RMSF within the connecting loop is indicative of decreased flexibility either when the PBD is bound to a model phosphopeptide **(e)**, or in the unbound state **(f)**.

Our PRM data suggested that EYA1 may also dephosphorylate pY425. We therefore tested the effect of Y425F and a Y425F/Y445F double mutant on PLK1 activation. Interestingly, neither Y425F nor the Y425F/Y445F double mutant significantly altered T210 phosphorylation. Given that both EYA4 and EYA1 support the activation of PLK1, this suggests that Y425 may only be a minor target of EYA1.

We then evaluated the ability of the Y445F mutant to rescue mitotic arrest caused by depletion of endogenous PLK1. Overexpression of WT PLK1 following depletion of endogenous PLK1 using a 3’UTR directed siRNA did not result in a detectable rescue of the mitotic arrest phenotype (Figure 5B). However, overexpression of Y445F PLK1 dramatically reduced the mitotic arrest phenotype triggered by endogenous PLK1 depletion, suggesting that the hyperactive Y445F mutant also has enhanced functionality (Figure 5B).

### Y445 phosphorylation inhibits the interaction with PLK1 activation complexes

Phosphorylation of PLK1 at pT210 by AURKA requires direct interaction between the PBD of PLK1 and a PDS on one of two AURKA cofactors: BORA in the cytoplasm, or CEP192 at centrosomes ^30^. As Y445 falls within the PBD of PLK1, and Y445F is hyperactive, we explored whether Y445 phosphorylation impacts the ability of PLK1 to interact with AURKA cofactors. We found that the Y445F mutant yielded a much stronger interaction with BORA compared to WT PLK1 in G2 and mitotically arrested cells (Figure 5C). The interaction between Y445F and CEP192 was also increased relative to WT PLK1 specifically in mitotic cells (Figure 5C). We observed a modest increase in the interaction between Y445F and AURKA itself in mitotic cells despite the transient nature of the PLK1-AURKA interaction (Figure 5C). Interestingly, in mitotic cells, Y445F also interacted more strongly than WT PLK1 with BUB1, a major kinetochore receptor for PLK1 in mitosis that interacts with the PBD of PLK1 ^49, 50^(Figure 5C). These data suggest that Y445 phosphorylation reduces the affinity of the PLK1 PBD for PLK1 interaction partners, which underpins a reduction in PLK1 activity and functionality.

### Phosphorylation of Y445 reduces flexibility within the PBD connecting loop and reduces substrate binding

We next investigated whether Y445 phosphorylation disrupts the structure and function of the PLK1 PBD. We performed molecular dynamics simulations using a published structure of the PBD of PLK1 bound to a model phosphopeptide (PDB: 1UMW)^32^. The binding free energy of the interaction was computed for the WT PLK1 PBD and PBD structures in which we simulated the phosphorylation of either Y445 or Y425. While phosphorylation of Y425 did not alter the computed binding free energy, phosphorylation of Y445 caused a decrease in the binding free energy (p < 0.05, Table S2). Additionally, pY445 is predicted to interact with R507 on the backside of the phosphopeptide binding pocket, causing a dramatic loss of flexibility in residues 502-506, which comprise a portion of the connecting loop between the two PBD domains that normally exhibit high flexibility (Figure 5D-F). Flexibility loss occurred irrespective of whether the PBD structure was bound to the model phosphopeptide (Figure 5D-F). These results suggest that phosphorylation of Y445 causes PBD dysfunction by perturbing both substrate binding and the overall structural dynamics of the PBD.

## Discussion

The EYA family is a biochemically unique group of protein tyrosine phosphatases; however, little is known about the substrates of the EYAs, or the downstream cellular processes impacted by EYA mediated dephosphorylation. Using BioID proximity proteomics we provide the first comprehensive characterization of EYA4 binding partners and find that many EYA4 interactors are involved in the regulation of mitosis. Additionally, we identify PLK1, one of the master regulatory kinases of mitosis, as the first known EYA4 phosphatase substrate, and describe a novel signalling pathway in which EYA4 and EYA1 regulate PLK1 through the dephosphorylation of pY445 during G2 (Figure 6). Dephosphorylation of PLK1 by the EYAs supports PLK1 activation, centrosomal maturation/separation, the prevention of spindle defects, and mitotic progression. Y445 on PLK1 has previously been shown to be phosphorylated by c-ABL, however this is the first report of its dephosphorylation and functional significance^51^.

**Fig. 6.**
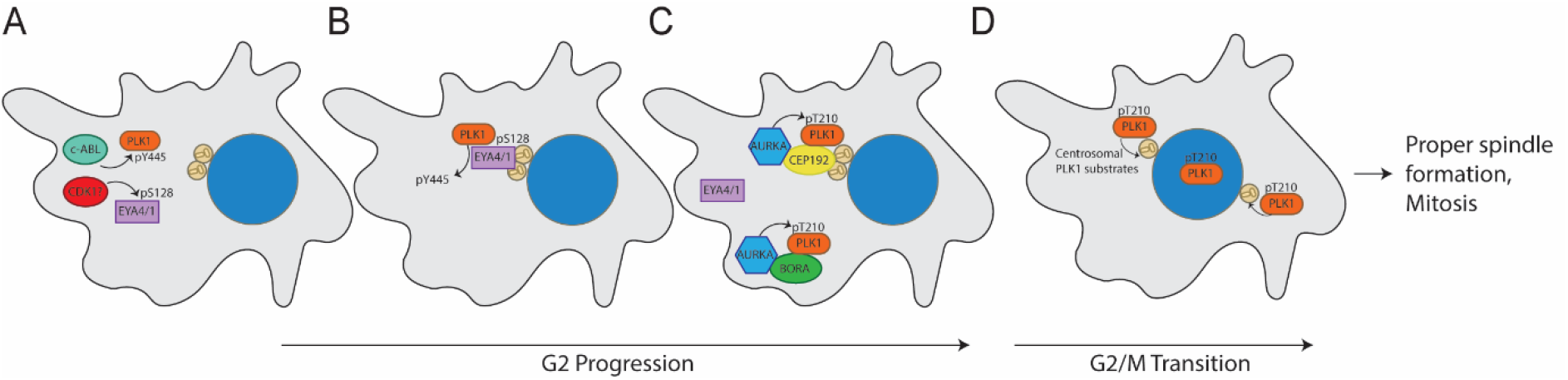
**a-d,** Model depicting dephosphorylation mediated regulation of PLK1 by EYA4/EYA1. **a,** Phosphorylation of PLK1 at Y445 by c-ABL and phosphorylation of EYA4/EYA1 within a PDS domain (S128 on EYA4: likely a CDK site) occurs during G2. **b,** EYA4/EYA1 interacts with PLK1 through an active PDS at centrosomes in G2 and dephosphorylates it at pY445. **c-d,** Dephosphorylated PLK1 has enhanced interactions with PLK1 activation complexes at centrosomes and in the cytoplasm, leading to enhanced pT210 phosphorylation and the promotion of PLK1 functions in centrosome maturation/separation, formation of the mitotic spindle, and mitotic progression.

The magnitude by which the individual EYAs support PLK1 activation is cell line dependent. This may be explained by variability in EYA protein levels (Figure S2B); however, functional differences between EYA4 and EYA1 also exist. Specifically, EYA4 dephosphorylates PLK1 at pY445 exclusively, while EYA1 appears to dephosphorylate both pY445 and pY425. Interestingly, the activating effect of an unphosphorylatable Y445F mutation was reduced by a concomitant unphosphorylatable Y425F mutation. This suggests that the degree to which EYA1 can promote PLK1 activity through pY445 dephosphorylation may in part depend on the initial phosphorylation level of pY425.

While the overall conclusions are supported by the use of an EYA phosphatase inhibitor, there were some phenotypic differences observed when compared to genetic depletion of the EYAs. In particular, treatment with benzarone reduced T210 phosphorylation as well as PLK1 levels in mitotic cells, compared to an exclusive reduction of T210 phosphorylation following knock down. EYA1 depletion also caused a slight increase in PLK1 levels in HeLa cells. Further, while both knock downs and benzarone treatment caused an induction of mitotic spindle defects, the magnitude of the induction was greater with benzarone treatment. Overall, these differences are likely explained by cellular adaptations that can occur during the course of a 72-hour siRNA time-course that do not have time to occur during the shorter benzarone treatment. However, it is also possible that the transcriptional activity of the EYAs can modulate some of their effects on PLK1 and mitosis.

Mechanistically, the dephosphorylation of pY445 promotes PLK1 activation in mitosis by supporting the interaction between PLK1 and PLK1-activation complexes. Phosphorylation of Y445 is predicted to decrease the interaction of the PLK1 PBD with a model peptide, as well as dramatically decrease the flexibility within the connecting loop of the PLK1 PBD structure. We propose that these changes have an additive effect, which results in the loss of PBD-dependent interactions. Given that the predicted reduction in binding between the PBD and model peptide is relatively modest, we expect that the loss of connecting loop flexibility is particularly important for *in-vivo* interactions. However, this is difficult to directly assess, as mutations in the connecting loop that experimentally alter flexibility would almost certainly have other intra and intermolecular effects that could not be controlled for.

PLK1 was one of 17 centrosomal EYA4 binding partners identified by BioID, which suggests that EYA4 may have additional centrosomal functions. A recent paper reported that EYA2 can localize to and support the function of centrosomes, even though it does not have an obvious PDS sequence ^17^. While it is possible that EYA2 has unrelated regulatory functions at centrosomes, our BioID data, as well as previous observations, suggest that both EYA2 and EYA1 can interact with EYA4. This raises the possibility that the EYAs may function in a complex, thereby enabling EYA2 to contribute to the dephosphorylation of PLK1 ^52^. These results highlight the need for further characterization of the roles of EYA proteins in centrosome biology.

Overall, we have characterized a new EYA-PLK1 signalling pathway with important implications for the structural regulation of PLK1, centrosome biology, and mitosis. Inactivation of the EYA-PLK1 signalling pathway potently induces mitotic cell death. This work provides a novel mechanism by which EYA phosphatase inhibitors may illicit cell killing in a therapeutic context, and justifies further testing of EYA phosphatase inhibitors in preclinical tumour models.

## Methods

### Vectors

All expression constructs were cloned into PCMV6 expression vectors. EYA4 and PLK1 mutants were generated using gBlocks™ Gene Fragments and restriction cloning.

### Cell culture, transfection, and gene knockdown

Cells were maintained in DMEM with 10% FCS in a sterile humidified incubator at 37°C with 10% CO_2_ and 20% O_2_. All cell lines were mycoplasma free and verified by STR profiling through CellBank Australia. Cells were passaged and harvested with trypsin. Cells were transfected at approximately 50% confluency with plasmid DNA using FuGENE® 6 or FuGENE® HD at a 3:1 ratio of FuGENE to DNA, according to manufacturer recommendations (Promega). For transient transfections plasmid DNA expression was allowed to proceed for 48 hours prior to downstream analysis. Gene knockdown was performed using Lipofectamine™ RNAiMAX according to manufacturer instructions (Thermo Fisher Scientific). Media containing RNAiMax and siRNA was changed, or cells were split into fresh media, 24 hours post transfection. Unless otherwise stated, downstream analysis occurred 72 hours after initial siRNA transfection.

### Cell synchronization and drug treatments

For enrichment of cells in mitosis or G2, cells were first synchronized in S-phase for 18 hours using 2 mM thymidine. Thymidine containing media was then removed and cells were washed 4 times with fresh media. For arrest in mitosis, nocodazole (0.1ug/ml) was added to cells for 12 hours prior to harvest. For arrest in G2, RO-3306 was added to media 4 hours after release from thymidine at a concentration of 10μM and cells were incubated for 6 hours prior to harvest. RO-3306 arrested cells were released into mitosis by removal of RO-3306 containing media and 3 washes with fresh media. Benzarone was diluted in DMSO and added to cells at concentrations as listed in the text.

### Immunoprecipitation/Co-immunoprecipitation

Cells were lysed in co-immunoprecipitation buffer (20 mM HEPES-KOH pH 7.9, 200 mM NaCl, 10% (v/v) glycerol, 0.1% Triton X-100, 1 mM DTT) supplemented with cOmplete Mini EDTA-free protease inhibitor cocktail and PhoshoStop phosphatase inhibitor tabs (Roche) at 4 °C. Lysates were cleared by centrifugation at 16,000 x g for 25 minutes at 4°C. Simultaneously, 4 or 5 ug of appropriate antibodies or IgG control proteins were bound to protein G Dynabeads™ as per manufacturer instructions (Life Technologies). Equal portions of cleared lysates were added to antibody or IgG bound Dynabeads™ and incubated for 3 hours or overnight at 4°C. Beads were washed with co-immunoprecipitation buffer at 4°C 4x for co-immunoprecipitation and 6x for immunoprecipitation, followed by elution with a 2:1 mixture of 50 mM Glycine, pH 2.8 and 1x NuPAGE™ LDS sample buffer (Life Technologies) at 70°C for 10 minutes.

### BioID pulldowns

BioID-Myc or BioID-Myc-EYA4 plasmids were transfected into T150 flasks of 293Ts at 50% confluence in quadruplicate. After 24 hours, media was removed and replaced with media containing 50 μM biotin. After 24 hours incubation with biotin, cells were trypsinised, pelleted and resuspended in 500 μL GdmCL lysis buffer (6 M GdmCl, 100mM Tris pH8.5, 10mM TCEP, 40mM Iodoacetamide). Lysates were heated at 95°C for 5 minutes and then cooled on ice for 15 min. Samples were sonicated using a Bioruptor ® sonicator at for 5 duty cycles for 30 seconds on and 30 seconds off at 4°C. Lysates were then heated again at 95°C for 5 minutes and then cooled on ice for 15 min, followed by a 30 minute RT incubation in the dark to allow for iodoacetamide to react with Cys residues. Lysates were then diluted 1:1 with milliQ water, and protein was precipitated overnight at −20°C using 4 volumes of acetone. The following day the proteins were pelleted by centrifugation for 5 minutes at 1500 RPM, washed once with 80% acetone, and resuspended by sonication as before. Proteins were again pelleted by centrifugation, acetone aspirated and left to dry for 15 minutes. Dried pellets were resuspended by sonication as before in resuspension buffer (2 M urea, 50 mM Sodium Pyrophosphate pH 8, 50 mM Ammonium Bicarbonate, 0.2% SDS). Protein concentration was determined by BCA assay. Next, 2 mg of each lysate was used in overnight immunoprecipitation reactions with 20 mg of streptavidin coated dynabeads. The following day, the beads were washed for 3×5 minutes with rotation in resuspension buffer. Next, 10% of the beads were transferred to a new tube and subject to elution of biotinylated proteins using 1X LDS at 80°C for 10 minutes for downstream western blot analysis. The remaining 90% of the beads were washed 3x with 100 mM Ammonium Bicarbonate and then resuspended in 100 μl of fresh 100 mM Ammonium Bicarbonate with 8% Acetonitrile (ACN) and 5 μg trypsin. On bead trypsin digestion was carried out overnight at 37°C shaking at 2000 RPM.

### C18 desalting of tryptic peptides

C18 stagetips were made using a syringe punch to pack to 2 layers of C18 filter into a low-retention p200 tip. C18 stagetips were activated by centrifuging through 100 μl of 100% acetonitrile and equilibrated with 0.1% formic acid (FA) and 0.1% trifluoroacetic acid (TFA). Tryptic peptides were prepared for desalting by the addition of 30 uL of 3.2 M KCL, 11 ul 150 mM KH_2_PO_4_ and 19 ul of 100% TFA followed by centrifugation at max speed for 15 minutes and re-tubing. Peptides were partially dried in a speedyvac for 30 minutes and then added to pre-equilibrated C18 stagetips and centrifuged at 1500 RPM. C18 bound peptides were washed twice with 100 μl of 0.1% FA and 0.1% TFA. Peptides were eluted with 40% ACN in 0.1% FA and fully dried in a speedyvac. Finally, samples were resuspended in 6 μl of mass spec buffer (1% FA, 2% ACN).

### LC-MS/MS and analysis for BioID and pY IPMS

Peptides were separated using a Dionex Ultimate 3000 UHPCL system with a nanoflow silica column packed with 1.9μM C18 beads using a 195-minute gradient. Mass spectrometry was performed using a Thermo Q Exactive instrument. The MS was operated in data-dependent mode with an MS1 resolution of 35,000, automatic gain control target of 3e6, and a maximum injection time of 20 milliseconds. The MS2 scans were run as a top 20 method with a resolution of 17,500, an automatic gain control target of 1e5, a maximum injection time of 25ms, a loop count of 20, an isolation window of 1.4 m/z and a normalized collision energy of 25.

Raw MS spectra data were matched, and label free quantification performed using MaxQuant. Default settings were used with the exception of LFQ min count, which was set to 1, matching from and to was enabled as well as matching between runs. Additionally, the match time was set to 1.5 min. For matching the following FASTA file was used: UP000005640_9606. Raw excel files were exported to Perseus where data was log2 transformed, potential contaminants removed, and imputation performed. P-values were generated in Perseus using a student’s T-test with a false discovery rate of 0.01. The presence of EYA4 and the established EYA4 interactors SIX1 and SIX2 among the high confidence EYA4 interactors in the BioID dataset further validated the approach.

For the analysis of BioID interactors with localization to mitotic structures pertaining to figure 1B we reduced the stringency for inclusion to include medium confidence EYA4 interactors (Log2 fold difference >1). Subcellular localization data was taken from uniprot as well as primary literature searches for each protein in combination with the search term “mitosis.”

### Identification and quantification of PLK tyrosine phosphorylated sites by mass spec

Myc-tagged PLK1 was overexpressed in triplicate or quadruplicate for each treatment in 293T cells. Treatments included siRNA against EYA4, EYA1 or a control sequence, overexpression of EYA4 Ydef or EYA4 wt, or incubation with benzarone (10uM) or DMSO for 2 hours prior to cell harvesting. Cells were pelleted, and PLK1 was immunopurified and eluted as described in the previous section. Immunopurified PLK1 was resolved on 4-12% Bis-Tris precast mini gels (Life Technologies). Following electrophoresis, gels were subjected to colloidal Coomassie staining according to the Anderson method ^53^. Bands corresponding to PLK1-Myc were excised, destained, reduced, alkylated, and digested in-gel with trypsin, following a previously described protocol ^54^. Peptides were fully dried down in a speedyvac and resuspended in 6 μl of mass spec buffer.

Peptides were separated using a Dionex Ultimate 3000 UHPCL system with a nanoflow silica column packed with 1.9μM C18 beads using a 130-minute gradient. Mass spectrometry was performed using a Thermo Q Exactive instrument operating in parallel reaction monitoring (PRM) mode using an inclusion list populated with phospho or unmodified mass and charge peptide data generated from pilot experiments (Table S2). MS2 scans were performed with a resolution of 17,500, an automatic gain control target of 5e4, a maximum injection time of 50 milliseconds, an isolation window of 1.6 m/z and a normalized collision energy of 30.

Skyline was used to extract chromatogram data and identify peaks from the PRM scans. The sum of product ion peak areas for individual peptides was quantified in accordance with the Skyline PRM tutorial. The ratio of phosphopeptide peak intensities was taken relative to corresponding unmodified peptides. Some datapoints were removed from analysis when precursor ion peaks were poorly identified, or in the case of clear outliers. Data presented in figure 2d is expressed as percent differences relative to control treatments and each datapoint represents at least biological duplicate.

### Raw MS data

The mass spectrometry proteomics data have been deposited to the ProteomeXchange Consortium via the PRIDE partner repository with the dataset identifiers PXD036405, PXD036430, and PXD037064.

### In-vitro phosphatase assay

The catalytic domain of human EYA4 (residues 367 – 639) was sub-cloned as a poly-His fusion construct with a TVMV cleavage site in the vector pDEST-527. Fusion protein was purified by Ni-NTA chromatography followed by ion exchange (Fast-Q) and size-exclusion chromatography over a Superdex-75 column. Phosphopeptides were synthesized by Lifetein. Peptide assays were conducted in 20 mM MES pH 6, 150 mM NaCl, 2 mM MgCl2, using a range of peptide concentrations (0 – 400 μM) and the Biomol assay. Data was analysed using non-linear regression in GraphPad PRISM assuming Michaelis-Menten kinetics.

### Immunoblotting

Western blot lysates were produced in RIPA buffer (Pierce #89900) supplemented with cOmplete Mini EDTA-free protease inhibitor cocktail and PhoshoStop phosphatase inhibitor tabs (Roche) and 1 mM DTT at 4 °C. Lysates were cleared by centrifugation at 16,000 x g for 25 minutes at 4 °C. Proteins were separated using 3-8% or 7% Tris-Acetate, or 4-12% Bis-Tris precast mini gels (Life Technologies) and transferred to PVDF membranes at 70V for 2 hours or overnight (Immobilon P, Millipore). Ponceau S staining was used to verify even transfer (Sigma Aldrich). Membranes were optionally cut to detect multiple proteins. Blocking and antibody incubations were then performed with either 5% non-fat milk or bovine serum albumin (fraction V) in PBST or TBST. Bands were visualized with HRP-conjugated secondary antibodies (DAKO) followed by application of a chemiluminescent reagent (Thermo Scientific). Stripping with Restore™ PLUS stripping buffer (Thermo Scientific) and reprobing was performed in some cases when the subsequent primary was of a different species; however, blots were never stripped more than once. Densitometry was performed in ImageJ on unaltered images.

### Immunofluorescence and Click-IT EdU staining

Cells were grown on coverslips pre-treated with Alcian blue stain to promote adherence (Sigma). For experiments involving EdU incorporation, EdU was added to the media for 1 hr prior to fixation at a final concentration of 10 μM. Cells were washed with PBS and then fixed using freshly prepared 4% paraformaldehyde (Sigma) for 10 minutes at RT followed by 2X PBS washes and then permeabilization with KCM buffer (120 mM KCl, 20 mM NaCl, 10 mM Tris pH 7.5, 0.1% Triton), or 0.2% Triton in PBS. If required, Click-It EdU staining was performed as per the manufacturer’s recommendations (Thermo Scientific). Next, blocking was performed using antibody-dilution buffer (20 mM Tris–HCl, pH 7.5, 2% (w/v) BSA, 0.2% (v/v) fish gelatin, 150 mM NaCl, 0.1% (v/v) Triton X-100 and 0.1% (w/v) sodium azide) for 1 hr at RT. Cells were incubated with primary antibodies for 1 hr at RT or overnight at 4°C followed by 3 × 10 minute washes in PBS. Cells were then incubated with Alexa Fluor conjugated secondary antibodies (Thermo Scientific) at a 1:500 or 1:750 dilution for 1 hr at RT. Cells were again washed for 3 × 10 minutes in PBS followed by staining in DAPI solution for 20 minutes (Sigma). Coverslips were mounted on slides in ProLong™ Gold antifade. Images were acquired with a Zeiss Axio Imager microscope.

### Imaging and Image Analysis

Fixed cell microscopy images were acquired with a Zeiss Axio Imager microscope. For automated image analysis images were converted from (.CZI) to (.TIFF) and imported into Cellprofiler v2.2 or v4.1 ^55^. Using custom image analysis pipelines we employed intensity-based thresholding strategies to mask individual objects (nuclei, centrosomes) and intensity-based measurements were made within these masks using the MeasureObjectIntensity module. Cytoplasmic area was approximated by expanding nuclear objects by 25 pixels in every direction. For analysis of centrosome number and intensity in prophase cells, images were imported into ImageJ. Prophase was staged by nuclear morphology and the positivity for H3 S10 phosphorylation and centrosomes were identified using the manual selection tool and intensity determined using the measure function. Mitotic defects were scored manually in image J.

### Live cell imaging and analysis using the Incucyte

For siRNA experiments knockdowns were performed in T75 flasks. Forty-eight hours later cells were reseeded into 24 well plates at 50% confluence and added to the incucyte. Imaging and data collection was performed 72 hours after initial knockdown. Overexpression experiments were performed directly within 24 well plates and imaging and data collection performed 48 hours after overexpression. Drug treatments were performed four hours before imaging and data collection. Phase contrast images were taken at 10X at fixed intervals for given experiments. Image analysis was performed manually.

### Molecular dynamics simulations

#### Preparation of structures

The crystal structure of the polo-box domain (PBD) of polo-like kinase 1 (Plk1) bound to a consensus phosphopeptide (PDB code 1UMW ^32^) was used as the initial structure for molecular dynamics (MD) simulations. The N- and C-termini of PLK1 PBD were capped by an acetyl group and N-methyl group, respectively. The protonation states of residues were determined by PDB2PQR. ^56^ In total, six PBD systems were set up: apo PBD, PBD complexed with phosphopeptide, apo PBD with Y445 phosphorylated (PBD-pY445), PBD-pY445 complexed with phosphopeptide, apo PBD with Y425 phosphorylated (PBD-pY425), and PBD-pY425 complexed with phosphopeptide. Each system was solvated with TIP3P water molecules ^57^ in a periodic truncated octahedron box such that its walls were at least 10 Å away from the protein, followed by charge neutralisation with sodium or chloride ions.

#### Molecular dynamics

Four independent MD simulations were carried out on each of the PBD systems. Energy minimisation and MD simulation were performed with the PMEMD module of AMBER 18 ^58^ using the ff14SB ^59^ force field. Parameters for phosphorylated residues were used as described by Homeyer *et al*. ^60^ A time step of 2 fs was used and the SHAKE algorithm ^61^ was implemented to constrain all bonds involving hydrogen atoms. The particle mesh Ewald method ^62^ was used to treat long-range electrostatic interactions under periodic boundary conditions. A cutoff distance of 9 Å was implemented for nonbonded interactions. The non-hydrogen atoms of the protein and peptide were kept fixed with a harmonic positional restraint of 2.0 kcal mol^−1^ Å^−2^ during the minimization and equilibration steps. Energy minimization was carried out using the steepest descent algorithm for 1000 steps, followed by another 1000 steps with the conjugate gradient algorithm. Gradual heating of the systems to 300 K was carried out at constant volume over 50 ps followed by equilibration at a constant pressure of 1 atm for another 50 ps. The restraints were removed for subsequent equilibration (2 ns) and production (300 ns for the WT PBD and PBD-pY445 systems, 500 ns for the PBD-pY425 systems) runs, which were carried out at 300 K and 1 atm, using a Langevin thermostat ^63^ with a collision frequency of 2 ps^−1^ and a Berendsen barostat ^64^ with a pressure relaxation time of 2 ps, respectively.

#### Binding free energy calculations

Binding free energies for the PBD–peptide complexes were calculated using the molecular mechanics/generalized Born surface area (MM/GBSA) method ^65^ implemented in AMBER 18.^58^ Two hundred equally-spaced snapshot structures were extracted from the last 100 ns of each of the trajectories, and their molecular mechanical energies calculated with the sander module. The 10 closest water molecules to the phosphothreonine in the phosphopeptide ligand in each snapshot were preserved and considered as part of the receptor for the MM/GBSA calculations. The polar contribution to the solvation free energy was calculated by the pbsa ^66^ program using the modified generalized Born (GB) model described by Onufriev *et al.* ^67^, with the solute dielectric constant set to 2 and the exterior dielectric constant set to 80. The nonpolar contribution was estimated from the solvent accessible surface area using the molsurf ^68^ program with γ = 0.005 kcal Å^−2^ and β = 0. The contribution of entropy was considered to be the same for all complexes and therefore ignored, as the receptors are structurally very similar and the ligand is unchanged. ^69^

### Statistical analysis

When not otherwise stated in the text p-values were generated in GraphPad Prism by students t-tests for comparisons involving two groups, or one-way ANOVAs for three or more groups. Tukey’s or Dunnett’s multiple comparisons tests were used with ANOVA to generate individual p-values when the mean of each group was compared to every other group or when the mean of each group was compared to the control group respectively.

## Extended Data

**Supp. Fig. 1.**
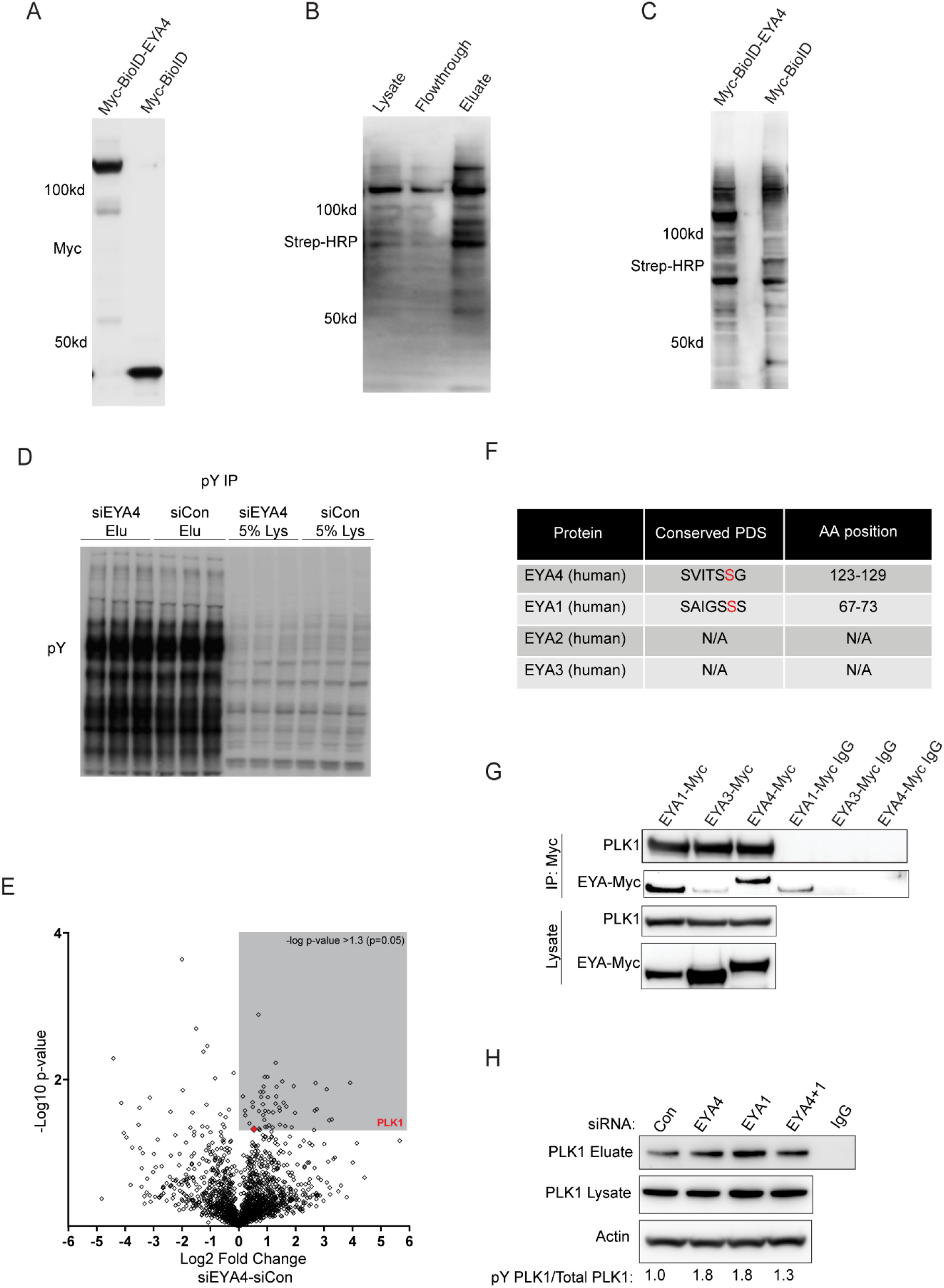
**a,** Western blot showing equal protein expression of EYA4-Myc-BioID and Myc-BioID constructs. **b,** Western blot showing enrichment of biotinylated proteins following precipitation with streptavidin. **c,** Western blot showing equal levels of biotinylated proteins in samples destined for trypsin digestion and mass spectrometry. **d,** Immunoprecipitation and western blotting of tyrosine phosphorylated proteins using an anti-phosphotyrosine antibody following depletion of EYA4 showing IP lysates (Lys) and Eluates (Elu). **e,** Volcano plot of immunoprecipitated tyrosine phosphorylated proteins identified by mass spectrometry, proteins with increased tyrosine phosphorylation following EYA4 depletion are on the right side (n=2). Grey coloured box represents 50 proteins with statistically significant increases in tyrosine phosphorylation (-log p-value > 1.3). PLK1 is indicated on the plot. **f,** Table showing conserved putative PDS and phosphosite in EYA4 and EYA1, but not present in EYA2 or EYA3. **g,** PLK1 coimmunopurification with Myc-tagged EYA4, and EYA1, but not EYA3, in G2 arrested cells. **h,** Immunoprecipitation of tyrosine phosphorylated proteins and western blotting. Densitometry values of PLK1 in the eluate (pY PLK1) relative to total PLK1 in the lysate is under the blot.

**Supp. Fig. 2.**
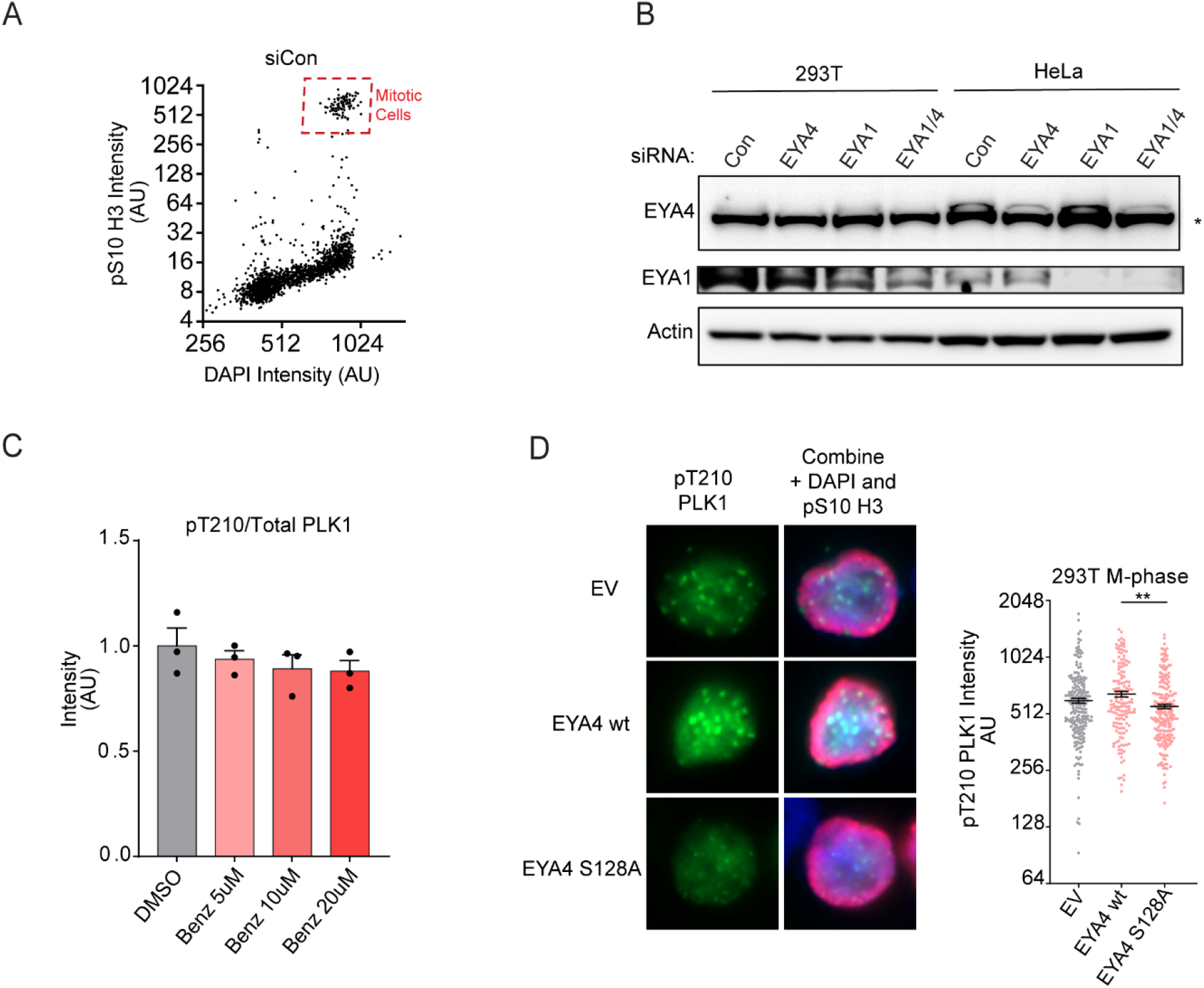
**a,** An example plot showing the identification of pS10 H3 positive and high DAPI (4N DNA content) mitotic cells (red box). **b,** Western blots confirming EYA4, EYA1 and combination knockdowns in asynchronous 293T and HeLa cells. Asterisk represents a non-specific band on the EYA4 blot. **c,** Western blot quantitation related to figure 3C. The ratio of pT210 PLK1 to total PLK1 in mitotic HeLa cells was reduced dose dependently by increasing concentrations of benzarone (ns). **d,** Mitotic 293T cells were identified by positive staining for pS10 H3 in high throughput image analysis experiments. Staining and quantitation were performed for pT210 phosphorylation of PLK1 in unsynchronized mitotic cells following overexpression of EYA4 or EYA4 S128A (p < 0.01).

**Supp. Fig. 3.**
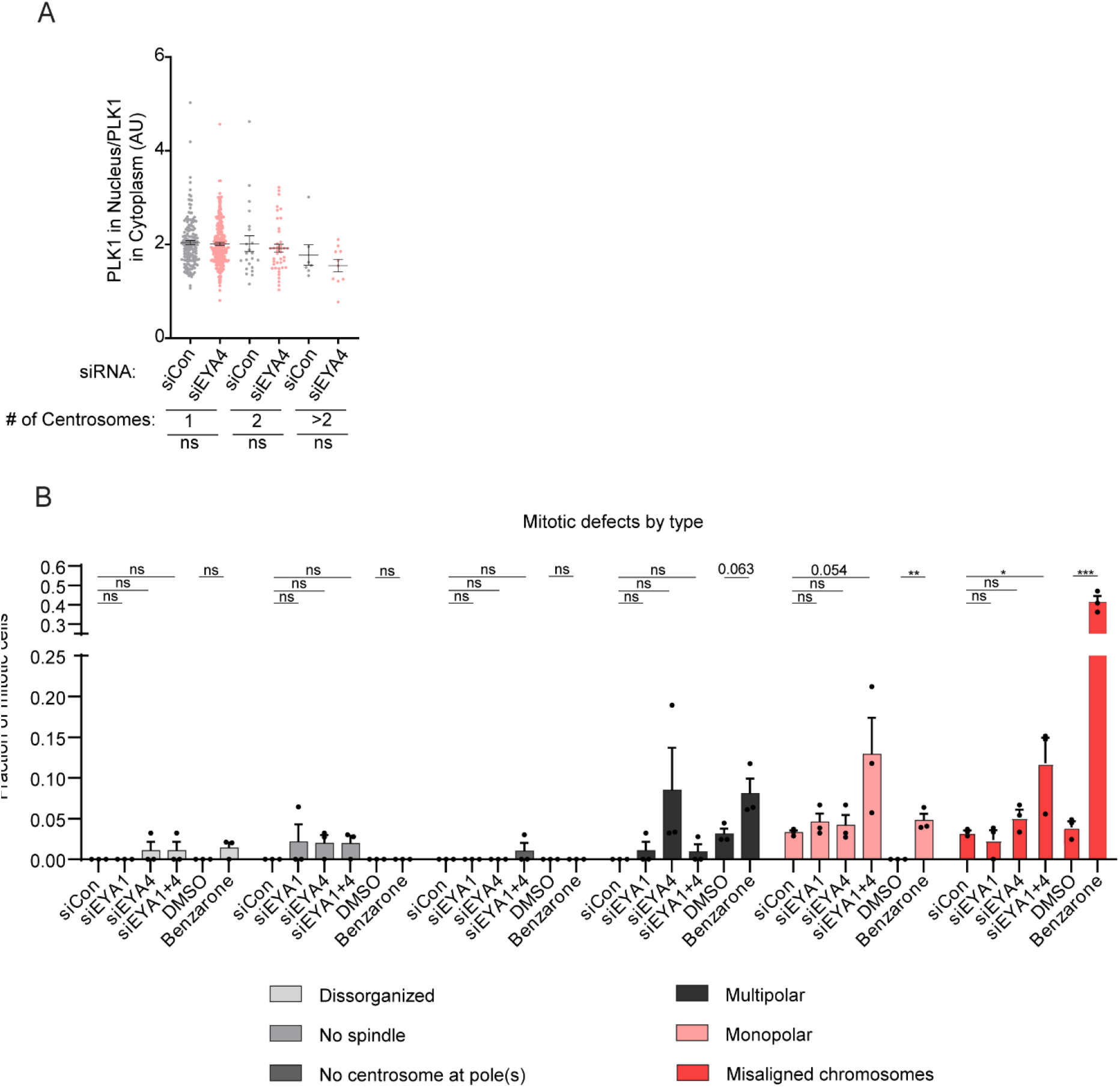
**a,** Immunofluorescence quantitation related to figure 3D-F. The ratio of PLK1 in the nucleus relative to that in the cytoplasm was unchanged in unsynchronized G2 cells following EYA4 depletion and irrespective of centrosome number. **b,** Spindle defect subdivision by type related to figure 3G-H. Co-depletion of EYA4 and EYA1 induced a moderate number of monopolar spindles as well as spindles with misaligned chromosomes (p = 0.054 and < 0.05, respectively). Treatment with benzarone induced a moderate amount of multipolar and monopolar spindles (p = 0.063 and < 0.01 respectively). Additionally, benzarone caused most spindles to have misaligned chromosomes (p < 0.001).

**Supp. Fig. 4.**
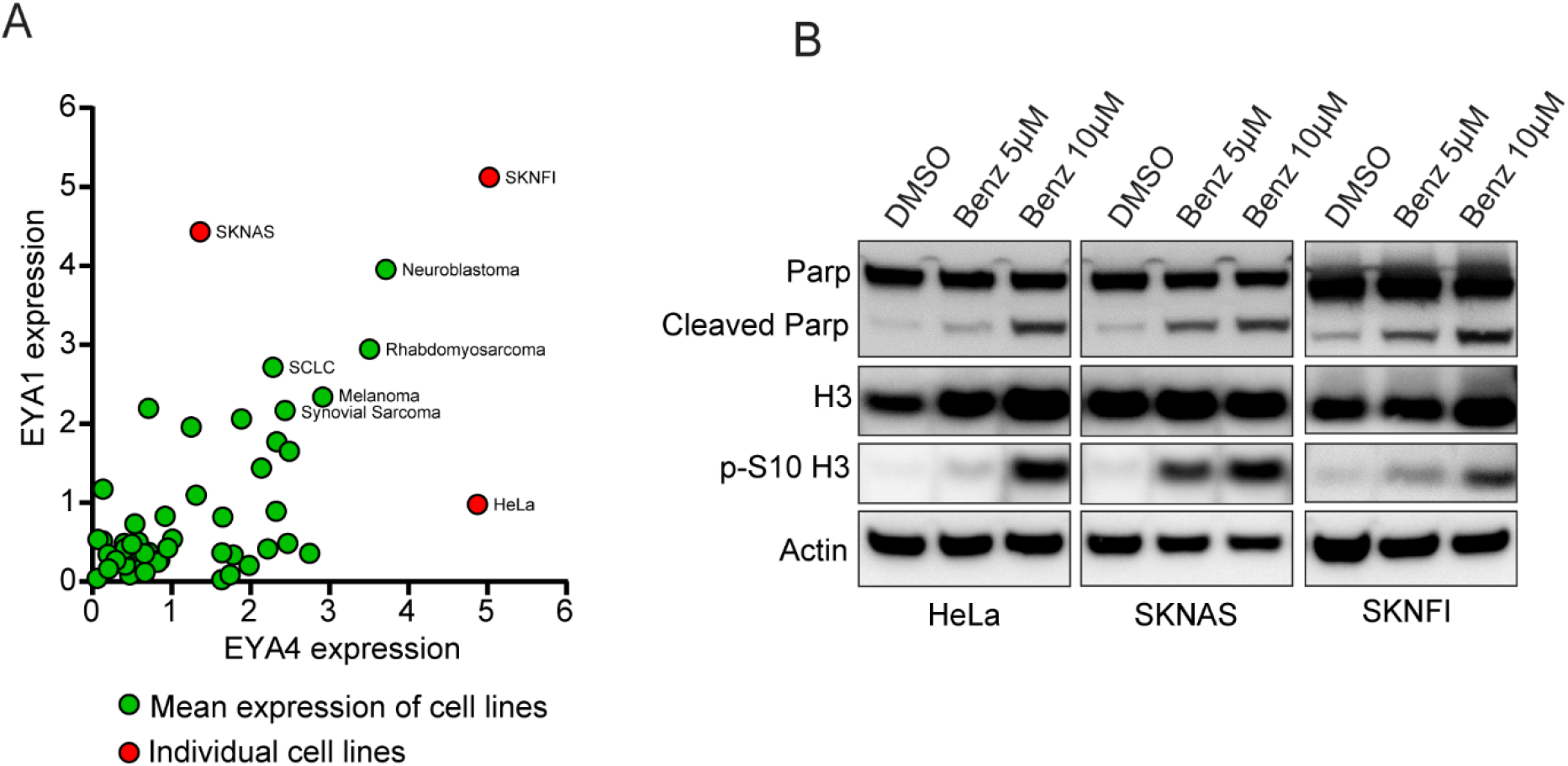
**a,** Data from a public gene expression database (DepMap Portal) indicated that HeLa cells had comparatively high expression of EYA4, but relatively low expression of EYA1. Therefore, we identified cells with high levels of EYA1 but low EYA4 (SKNAS) and high levels of both EYA4 and EYA1 (SKNFI) to test the relative contribution of each gene to sensitivity to benzarone. **b,** Western blot of cleaved PARP and pS10 H3 in response to treatment with benzarone.

**Supp. Fig. 5.**
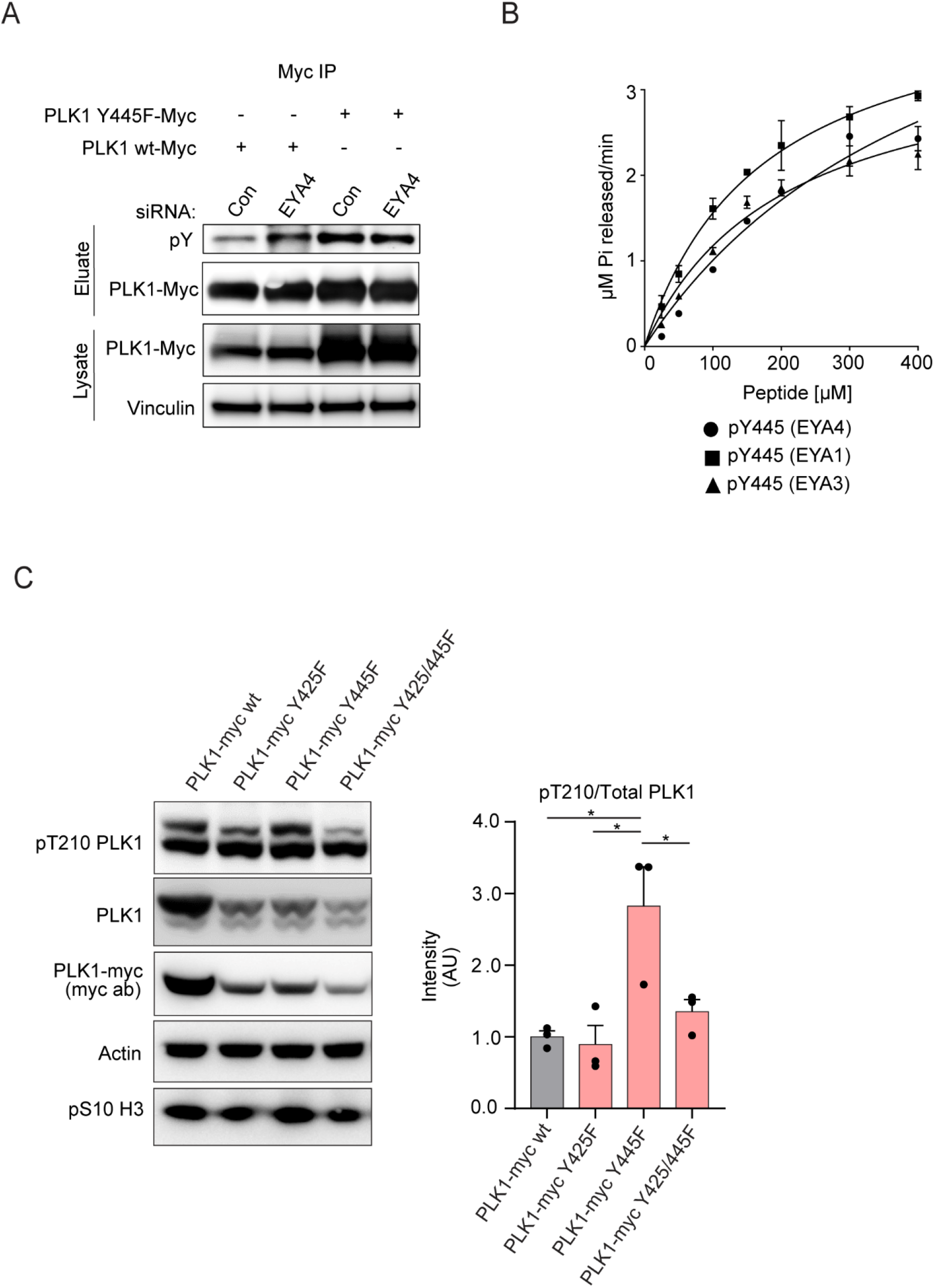
**a,** Immunoblotting of total tyrosine phosphorylation on immunoprecipitated PLK1 or a Y445F mutant in cells arrested in G2 with RO3306 following treatment with a control siRNA or one targeting EYA4.. **b,** *In-vitro* phosphatase assays using the phosphatase domains of EYA4, EYA1, and EYA3 and a pY445 phosphopeptide. All dephosphorylation reactions fit with Michaelis-Menten kinetics, error bars represent standard deviations. **c,** Expression of myc-tagged WT PLK1 or phosphosite mutants including Y425F, Y445F and a combination mutant in nocodazole arrested mitotic 293T cells. Densitometry of the T210 PLK1/total PLK1 ratio is presented (p < 0.05).

## Notes

### Competing Interest Statement

The authors have declared no competing interest.

## References

1. Kong, D. et al. EYA1 promotes cell migration and tumor metastasis in hepatocellular carcinoma. Am J Transl Res 11, 2328–2338 (2019).

2. Xie, P. et al. The deubiquitinase OTUB1 fosters papillary thyroid carcinoma growth through EYA1 stabilization. J Cell Mol Med 25, 10980–10989 (2021).

3. Kim, J. et al. Phage display targeting identifies EYA1 as a regulator of glioblastoma stem cell maintenance and proliferation. Stem Cells 39, 853–865 (2021).

4. Xu, H., Jiao, Y., Yi, M., Zhao, W. & Wu, K. EYA2 Correlates With Clinico-Pathological Features of Breast Cancer, Promotes Tumor Proliferation, and Predicts Poor Survival. Front Oncol 9, 26 (2019).

5. Ren, L., Guo, D., Wan, X. & Qu, R. EYA2 upregulates miR-93 to promote tumorigenesis of breast cancer by targeting and inhibiting the STING signaling pathway. Carcinogenesis, bgab001 (2021).

6. Xu, S., Lian, Z., Zhang, S., Xu, Y. & Zhang, H. CircGNG4 Promotes the Progression of Prostate Cancer by Sponging miR-223 to Enhance EYA3/c-myc Expression. Front Cell Dev Biol 9, 684125 (2021).

7. Miller, S.J. et al. Inhibition of Eyes Absent Homolog 4 expression induces malignant peripheral nerve sheath tumor necrosis. Oncogene 29, 368–379 (2010).

8. Yang, J. & Du, X. Genomic and molecular aberrations in malignant peripheral nerve sheath tumor and their roles in personalized target therapy. Surgical Oncology 22, e53–e57 (2013).

9. Zhu, J., Hu, L.-B., Zhao, Y.-P. & Zhang, Y.-Q. Prognostic Role of EYA4 in Lower Grade Glioma with IDH1 Mutation and 1p19q Co-Deletion. World Neurosurg 149, e1174–e1179 (2021).

10. Kong, D. et al. SIX1 Activates STAT3 Signaling to Promote the Proliferation of Thyroid Carcinoma via EYA1. Front Oncol 9, 1450 (2019).

11. Li, Z., Qiu, R., Qiu, X. & Tian, T. EYA2 promotes lung cancer cell proliferation by downregulating the expression of PTEN. Oncotarget 8, 110837–110848 (2017).

12. Li, Z., Qiu, R., Qiu, X. & Tian, T. EYA4 Promotes Cell Proliferation Through Downregulation of p27Kip1 in Glioma. Cell Physiol Biochem 49, 1856–1869 (2018).

13. Pandey, R.N. et al. The Eyes Absent phosphatase-transactivator proteins promote proliferation, transformation, migration, and invasion of tumor cells. Oncogene 29, 3715–3722 (2010).

14. Wang, Y. et al. The Protein Tyrosine Phosphatase Activity of Eyes Absent Contributes to Tumor Angiogenesis and Tumor Growth. Mol Cancer Ther 17, 1659–1669 (2018).

15. Wang, Y. et al. Targeting EYA3 in Ewing Sarcoma Retards Tumor Growth and Angiogenesis. Mol Cancer Ther 20, 803–815 (2021).

16. Wu, K. et al. EYA1 phosphatase function is essential to drive breast cancer cell proliferation through cyclin D1. Cancer Res 73, 4488–4499 (2013).

17. Zhang, G. et al. Targeting EYA2 tyrosine phosphatase activity in glioblastoma stem cells induces mitotic catastrophe. J Exp Med 218, e20202669 (2021).

18. Eisner, A. et al. The Eya1 Phosphatase Promotes Shh Signaling during Hindbrain Development and Oncogenesis. Developmental Cell 33, 22–35 (2015).

19. Yuan, B. et al. A phosphotyrosine switch determines the antitumor activity of ERβ. J Clin Invest 124, 3378–3390 (2014).

20. Krueger, A.B. et al. Allosteric inhibitors of the Eya2 phosphatase are selective and inhibit Eya2-mediated cell migration. J Biol Chem 289, 16349–16361 (2014).

21. Tonks, N.K. Protein tyrosine phosphatases – from housekeeping enzymes to master regulators of signal transduction. The FEBS Journal 280, 346–378 (2013).

22. He, R.-j., Yu, Z.-h., Zhang, R.-y. & Zhang, Z.-y. Protein tyrosine phosphatases as potential therapeutic targets. Acta Pharmacol Sin 35, 1227–1246 (2014).

23. Stanford, S.M. & Bottini, N. Targeting Tyrosine Phosphatases: Time to End the Stigma. Trends in Pharmacological Sciences 38, 524–540 (2017).

24. Anantharajan, J. et al. Structural and Functional Analyses of an Allosteric EYA2 Phosphatase Inhibitor That Has On-Target Effects in Human Lung Cancer Cells. Mol Cancer Ther 18, 1484–1496 (2019).

25. Tadjuidje, E. et al. The EYA tyrosine phosphatase activity is pro-angiogenic and is inhibited by benzbromarone. PLOS ONE 7, e34806 (2012).

26. Krueger, A.B. et al. Identification of a selective small-molecule inhibitor series targeting the eyes absent 2 (Eya2) phosphatase activity. J Biomol Screen 18, 85–96 (2013).

27. Pandey, R.N. et al. Structure-activity relationships of benzbromarone metabolites and derivatives as EYA inhibitory anti-angiogenic agents. PLOS ONE 8, e84582 (2013).

28. Kim, D.I. et al. An improved smaller biotin ligase for BioID proximity labeling. MBoC 27, 1188–1196 (2016).

29. Roux, K.J., Kim, D.I., Raida, M. & Burke, B. A promiscuous biotin ligase fusion protein identifies proximal and interacting proteins in mammalian cells. Journal of Cell Biology 196, 801–810 (2012).

30. Joukov, V. & Nicolo, A.D. Aurora-PLK1 cascades as key signaling modules in the regulation of mitosis. Science Signaling (2018).

31. Schmucker, S. & Sumara, I. Molecular dynamics of PLK1 during mitosis. Mol Cell Oncol 1, e954507 (2014).

32. Elia, A.E.H. et al. The molecular basis for phosphodependent substrate targeting and regulation of Plks by the Polo-box domain. Cell 115, 83–95 (2003).

33. Elia, A.E.H., Cantley, L.C. & Yaffe, M.B. Proteomic Screen Finds pSer/pThr-Binding Domain Localizing Plk1 to Mitotic Substrates. Science 299, 1228–1231 (2003).

34. Kettenbach, A.N. et al. Quantitative Phosphoproteomics Identifies Substrates and Functional Modules of Aurora and Polo-Like Kinase Activities in Mitotic Cells. Science Signaling 4, rs5–rs5 (2011).

35. Lemmens, B. et al. DNA Replication Determines Timing of Mitosis by Restricting CDK1 and PLK1 Activation. Mol. Cell 71, 117–128.e113 (2018).

36. Gheghiani, L., Loew, D., Lombard, B., Mansfeld, J. & Gavet, O. PLK1 Activation in Late G2 Sets Up Commitment to Mitosis. Cell Rep 19, 2060–2073 (2017).

37. Macůrek, L. et al. Polo-like kinase-1 is activated by aurora A to promote checkpoint recovery. Nature 455, 119–123 (2008).

38. Bruinsma, W. et al. Spatial Separation of Plk1 Phosphorylation and Activity. Front Oncol 5 (2015).

39. García-Álvarez, B., Cárcer, G.d., Ibañez, S., Bragado-Nilsson, E. & Montoya, G. Molecular and structural basis of polo-like kinase 1 substrate recognition: Implications in centrosomal localization. PNAS 104, 3107–3112 (2007).

40. Reindl, W., Yuan, J., Krämer, A., Strebhardt, K. & Berg, T. Inhibition of Polo-like Kinase 1 by Blocking Polo-Box Domain-Dependent Protein-Protein Interactions. Chemistry & Biology 15, 459–466 (2008).

41. Lane, H.A. & Nigg, E.A. Antibody microinjection reveals an essential role for human polo-like kinase 1 (Plk1) in the functional maturation of mitotic centrosomes. Journal of Cell Biology 135, 1701–1713 (1996).

42. Lee, K. & Rhee, K. PLK1 phosphorylation of pericentrin initiates centrosome maturation at the onset of mitosis. Journal of Cell Biology 195, 1093–1101 (2011).

43. Mahen, R., Jeyasekharan, A.D., Barry, N.P. & Venkitaraman, A.R. Continuous polo-like kinase 1 activity regulates diffusion to maintain centrosome self-organization during mitosis. PNAS 108, 9310–9315 (2011).

44. Addis Jones, O., Tiwari, A., Olukoga, T., Herbert, A. & Chan, K.-L. PLK1 facilitates chromosome biorientation by suppressing centromere disintegration driven by BLM-mediated unwinding and spindle pulling. Nature Communications 10, 2861 (2019).

45. Ehlén, Å. et al. Proper chromosome alignment depends on BRCA2 phosphorylation by PLK1. Nature Communications 11, 1819 (2020).

46. Lénárt, P. et al. The Small-Molecule Inhibitor BI 2536 Reveals Novel Insights into Mitotic Roles of Polo-like Kinase 1. Current Biology 17, 304–315 (2007).

47. Vugt, M.A.T.M.v. et al. Polo-like Kinase-1 Is Required for Bipolar Spindle Formation but Is Dispensable for Anaphase Promoting Complex/Cdc20 Activation and Initiation of Cytokinesis *. Journal of Biological Chemistry 279, 36841–36854 (2004).

48. Krishnan, N. et al. Dephosphorylation of the C-terminal Tyrosyl Residue of the DNA Damage-related Histone H2A.X Is Mediated by the Protein Phosphatase Eyes Absent *. Journal of Biological Chemistry 284, 16066–16070 (2009).

49. Singh, P. et al. BUB1 and CENP-U, Primed by CDK1, Are the Main PLK1 Kinetochore Receptors in Mitosis. Mol. Cell 81, 67–87.e69 (2021).

50. Qi, W., Tang, Z. & Yu, H. Phosphorylation- and polo-box-dependent binding of Plk1 to Bub1 is required for the kinetochore localization of Plk1. MBoC 17, 3705–3716 (2006).

51. Yang, X. et al. Cervical Cancer Growth Is Regulated by a c-ABL–PLK1 Signaling Axis. Cancer Res 77, 1142–1154 (2017).

52. Kumar, P. et al. A Human Tyrosine Phosphatase Interactome Mapped by Proteomic Profiling. J Proteome Res 16, 2789–2801 (2017).

53. Dyballa, N. & Metzger, S. Fast and Sensitive Coomassie Staining in Quantitative Proteomics, in *Quantitative Methods in Proteomics*. (ed. K. Marcus) 47–59 (Humana Press, Totowa, NJ; 2012).

54. Breitkopf, S.B. & Asara, J.M. Determining in vivo phosphorylation sites using mass spectrometry. Curr Protoc Mol Biol Chapter 18, Unit18.19.11-27 (2012).

55. Carpenter, A.E. et al. CellProfiler: image analysis software for identifying and quantifying cell phenotypes. Genome Biology 7, R100 (2006).

56. Dolinsky, T.J. et al. PDB2PQR: expanding and upgrading automated preparation of biomolecular structures for molecular simulations. Nucleic Acids Research 35, W522–W525 (2007).

57. Jorgensen, W.L., Chandrasekhar, J., Madura, J.D., Impey, R.W. & Klein, M.L. Comparison of simple potential functions for simulating liquid water. Journal of Chemical Physics 79, 926–935 (1983).

58. Case, D.A. et al. (University of California, San Francisco, 2018).

59. Maier, J.A. et al. ff14SB: improving the accuracy of protein side chain and backbone parameters from ff99SB. Journal of Chemical Theory and Computation 11, 3696–3713 (2015).

60. Homeyer, N., Horn, A.H.C., Lanig, H. & Sticht, H. AMBER force-field parameters for phosphorylated amino acids in different protonation states: phosphoserine, phosphothreonine, phosphotyrosine, and phosphohistidine. Journal of Molecular Modeling 12, 281–289 (2006).

61. Ryckaert, J.P., Ciccotti, G. & Berendsen, H.J.C. Numerical integration of the Cartesian equations of motion of a system with constraints: molecular dynamics of n-alkanes. Journal of Computational Physics 23, 327–341 (1977).

62. Darden, T., York, D. & Pedersen, L. Particle mesh Ewald: an N•log(N) method for Ewald sums in large systems. Journal of Chemical Physics 98, 10089–10092 (1993).

63. Izaguirre, J.A., Catarello, D.P., Wozniak, J.M. & Skeel, R.D. Langevin stabilization of molecular dynamics. Journal of Chemical Physics 114, 2090–2098 (2001).

64. Berendsen, H.J.C., Postma, J.P.M., Vangunsteren, W.F., Dinola, A. & Haak, J.R. Molecular dynamics with coupling to an external bath. Journal of Chemical Physics 81, 3684–3690 (1984).

65. Srinivasan, J., Cheatham, T.E., Cieplak, P., Kollman, P.A. & Case, D.A. Continuum solvent studies of the stability of DNA, RNA, and phosphoramidate−DNA helices. Journal of the American Chemical Society 120, 9401–9409 (1998).

66. Luo, R., David, L. & Gilson, M.K. Accelerated Poisson-Boltzmann calculations for static and dynamic systems. Journal of Computational Chemistry 23, 1244–1253 (2002).

67. Onufriev, A., Bashford, D. & Case, D.A. Exploring protein native states and large-scale conformational changes with a modified generalized Born model. Proteins: Structure, Function & Bioinformatics 55, 383–394 (2004).

68. Connolly, M.L. Analytical molecular surface calculation. Journal of Applied Crystallography 16, 548–558 (1983).

69. Hou, T., Wang, J., Li, Y. & Wang, W. Assessing the performance of the MM/PBSA and MM/GBSA methods. 1. The accuracy of binding free energy calculations based on molecular dynamics simulations. Journal of Chemical Information and Modeling 51, 69–82 (2011).

